# DiADeM: differential analysis via dependency modelling of chromatin interactions with robust generalized linear models

**DOI:** 10.1101/654699

**Authors:** Rafał Zaborowski, Bartek Wilczyński

## Abstract

High throughput Chromosome Conformation Capture experiments have become the standard technique to assess the structure and dynamics of chromosomes in living cells. As any other sufficiently advanced biochemical technique, Hi-C datasets are complex and contain multiple documented biases, with the main ones being the non-uniform read coverage and the decay of contact coverage with genomic distance. Both of these effects have been studied and there are published methods that are able to normalize different Hi-C data to mitigate these biases to some extent. It is crucial that this is done properly, or otherwise the results of any comparative analysis of two or more Hi-C experiments are bound to be biased. In this paper we study both mentioned biases present in the Hi-C data and show that normalization techniques aimed at alleviating the coverage bias are at the same time exacerbating the problems with contact decay bias. We also postulate that it is possible to use generalized linear models to directly compare non-normalized data an that it is giving better results in identification of differential contacts between Hi-C matrices than using the normalized data.

## Introduction

Eukaryotic chromosomes are very tightly packed biopolymers in the space of the nucleus. Despite the ultra-dense packing, their biological functions such as the ability to replicate and to express genes, depends on reproducible structural dynamics [Rada-Iglesias et al., 2018]. Importantly, their structure is not necessarily completely reproduced between the cells in the same epigenetic state, however, certain contacts between parts of the chromosomes, such as the enhancer-promoter contacts or insulator-mediated loops are essential to ensure proper gene regulation [Eagen, 2018]. Chromosome Conformation Capture and related techniques have been developed in the last years in order to give the researchers the ability to measure the frequency of such contacts and make inferences about the regulatory relationships based on the relative contact frequencies [Dekker et al., 2002]. In particular, the development of the hi-throughput chromosome conformation capture technique (Hi-C) has allowed us to measure genome-wide statistical chromosome contact patterns from large populations of cells [Lieberman-Aiden et al., 2009]. The introduction of this technique a decade ago, and its subsequent improvements, gave us now access to multiple datasets containing measured average frequencies of contacts across wide variety of cell types in multiple species [Schmitt et al., 2016, Rudan et al., 2015].

While the importance of the Hi-C technique for studying chromosome structure and dynamics should not be underestimated, as with any sufficiently complex biochemical protocol, we need to take into account the natural biases of the method when analysing the data [Yaffe and Tanay, 2011]. Especially when one is attempting to make statements comparing two or more Hi-C datasets originating from different cell populations, one needs to be very cautious whether the inferred conclusions are not due to any intrinsic biases of the measurements itself. This is especially important in the case of Hi-C data, where due to the very high cost of obtaining the data, researchers are frequently facing the necessity to make comparative analyses without sufficient number of replicate experiments.

There are two very well documented biases in the Hi-C data: the non-uniform coverage, that needs to be removed if relative statements are to be made comparing contact frequencies between different regions in the same data; and the contact decay bias, that gives rise to the observed decrease in contact frequency of regions proportional to their distance in the linear space of the chromosome [Imakaev et al., 2012, Nagano et al., 2015]. The first bias is usually taken care of by the standard normalization techniques, however the second one is usually considered to be a biological phenomenon and not accounted for in the normalization. However, when comparing different Hi-C datasets one can clearly see that even in datasets coming from seemingly the same type of cells, significant differences in the decay rate can be observed between experiments, that cannot be explained by the biological differences between the studied cell populations. In such cases, when one is interested in identifying differential contacts between Hi-C maps, the results of the comparative analyses can be severely and systematically biased by the difference in decay rates.

In this paper we take on the problem of identifying and measuring the effects of both biases and whether the currently available normalization methods for Hi-C data are adequate for countering the observed biases in the data. We also propose a solution to the problems identified with current methods.

## Results

In this section we report the main results of our study. We start by describing the coverage and contact decay effects and explain how they affect differential analysis of Hi-C data. Then we show that existing Hi-C normalization methods fail to account for all issues resulting from differences in coverage and contact decay. Finally we present that raw Hi-C interaction patterns at given genomic distance are highly correlated even between different cell line datatsets. This observation enables to construct a linear background model allowing for discovery of local changes in contact intensity by testing for deviations from the expected pattern. Finally we introduce, a simple algorithm for detection of long range differentially interacting regions.

### Dominating signals in Hi-C data

There are 2 factors, which exhibit significant variation across Hi-C datasets: the coverage and contact decay. The variation is particularly large when samples were derived from different studies. However, significant shift in either of these two signals may be observed even during comparison of supposedly technical replicates from a single sample (see for instance IMR90 replicates from GSE63525 study - Supplemental Figure 1, 2) [Rao et al., 2014]. In perfect scenario we would like both datasets to have both coverage and contact decay invariable. This would allow to detect local differences in the number of interactions between pairs of chromatin regions, which is usually the purpose of Hi-C differential analysis. In absence of the above situation there is no straight-forward way for finding local contact differences. Below we demonstrate the influence of coverage and contact decay on Hi-C differential analysis.

#### Hi-C coverage bias

Like any sequencing based protocol an outcome of Hi-C experiment depends on the sequencing depth. The higher the number of reads, the larger signal to noise ratio may be achieved. However often reasearchers are forced to compare Hi-C datasets obtained from different studies, which usually implicates diverging coverages of corresponding contact maps (Figure 1 top panel). Because of this direct comparisons would be severely biased leading to enrichment of most cells in higher coverage matrix (Supplemental Figure 3). To avoid this problem contact matrices are usually normalized before conducting differential analysis. Among many normalization methods the most commonly used seems to be the Iterative Correction approach based on assumption of uniform visibillity of every region [Imakaev et al., 2012]. This method is largely effective in removing coverage bias (Figure 1 bottom panel). Unfortunately, as we point out in the next section such normalization may even amplify issues stemming from other biases during Hi-C differential analysis. Most other normalization methods are focusing on this effect ignoring the contact decay.

**Figure 1.**
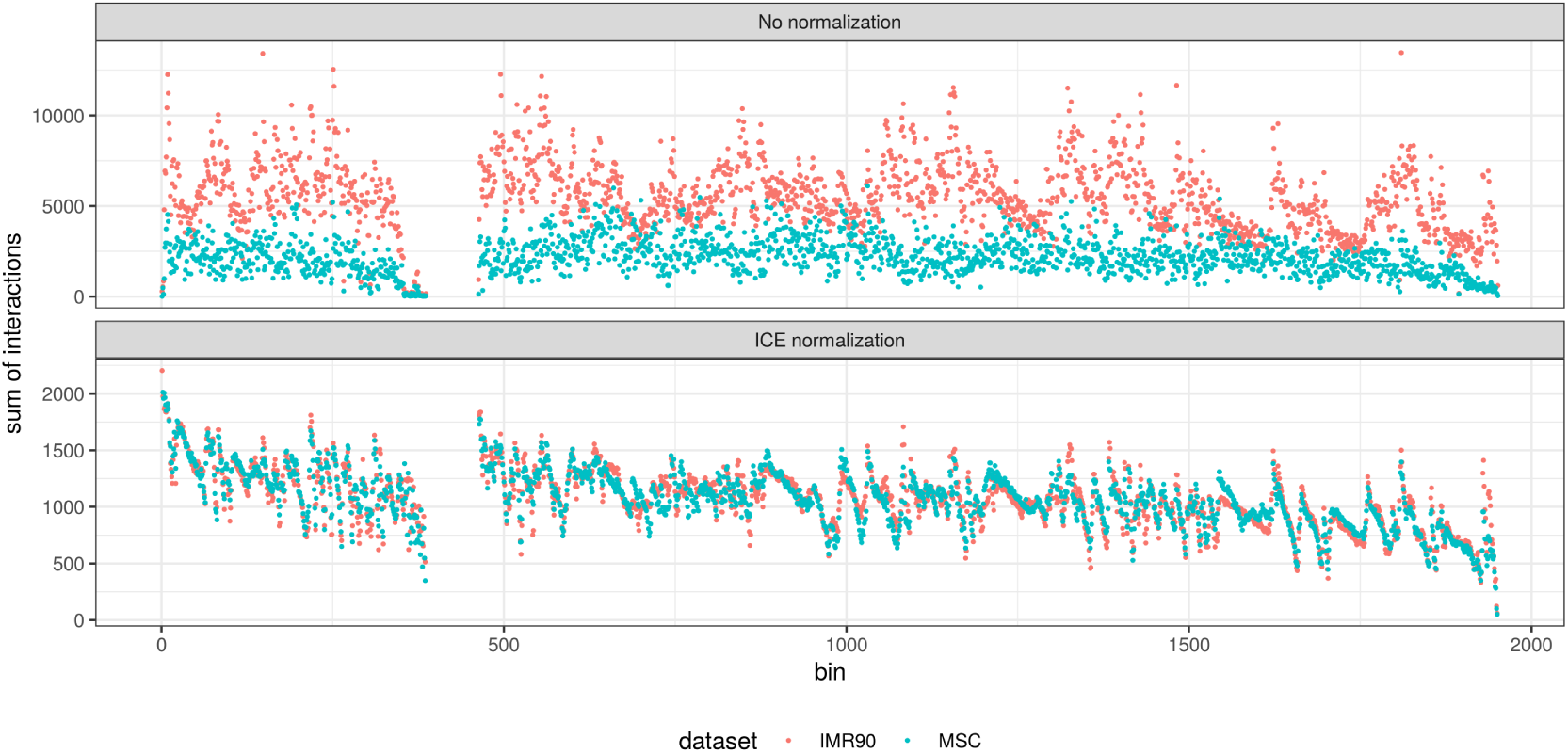
Coverages of human IMR90 and MSC cell Hi-C data, chromosome 18. Top panel - raw data. Bottom panel - ICE normalized.

#### Hi-C contact decay bias

The contact decay refers to the rate of change in number of interactions between a pair of chromatin regions as a function of their linear separation on the chromosome. Usually it is measured as the mean number of interactions between all pairs of loci at given distance (see Materials and methods for details). Intuitively this relationship reflects chromatin compaction and is very pronounced in Hi-C data (Figure 2 top). Such effect masks virtually any local differences allowing to only compare global compaction of chromatin. Iterative correction method mentioned in previous section is not particularly helpful this time as it does not remove contact decays difference (Figure 2 center and supplementary Figure 4). What is worse, different normalization methods may yield opposing patterns of relative contact decays between compared conditions (Figure 2 bottom and Supplemental Figure 4, 5). As can be seen in Figure 2 initial relationship between human IMR90 and MSC contact decays exhibit much stronger signal for IMR90 data set at almost entire range. Upon normalization this pattern changes significantly. Depending on which method is used, MSC contact intensity starts to exceed IMR90 at around 80th (HiCnorm) or 6th (ICE) diagonal [Hu et al., 2012]. Both these methods show consistent behaviour for higher ranges of contact decays (Figure2, third column, rows 2 and 3) however they diverge significantly at close genomic distances (Figure 2, second column, rows 2 and 3), where the most reliable interaction data is collected.

**Figure 2.**
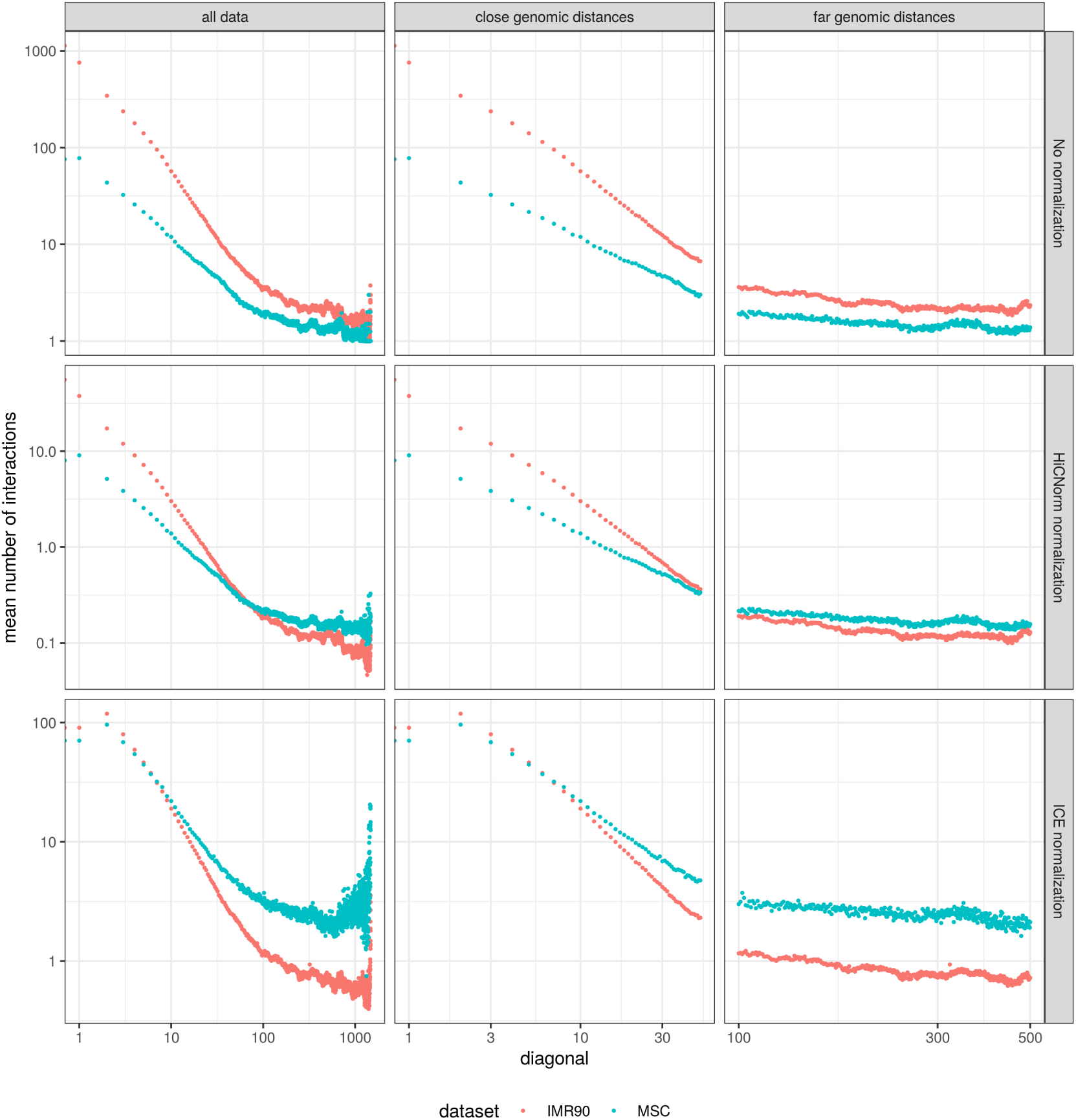
Contact decays of human IMR90 and MSC cell Hi-C data, chromosome 19. Top row - raw data, middle row - HiCnorm normalized, bottom row - ICE normalized.

### Correlations between raw contact intensities

When analyzing different Hi-C matrices, we discovered that relationship between raw interaction intensities of two Hi-C datasets at given loci separation exhibit approximately linear behaviour with non constant (funnel like) variance (Supplemental Figure 6, 7, 8). Despite different coverages, Pearson correlations between contact intensities are evident and statistically significant for pairs of regions separated by up to few megabases (Figure 3a top). Surprisingly this trend is not limited to replicates (Supplemental Figure 9, 10), but it is also pronounced between different study datasets, being much weaker, but still highly significant (Figure 3b bottom). The maximum separation of interacting regions, which preserves statistical significance between interaction intensities varies, typically ranging from 4 to 8 or even 16 Mbp. The precise value of separation threshold depends on the dataset quality (notably sequencing depth) and the chromosome size (Supplemental Table 1). Additionally we noticed that the distributions of paired interactions for certain ranges of genomic distances vary no more then what would be expected by chance (in terms of statistical distance - see Materials and methods for details). Using energy statistic and multivariate distribution equality test based on it we developed pooling procedure allowing to aggregate diagonals in larger groups (see Figure 3b Materials and methods for details). The main benefit of pooling is the increased number of observations for some training sets (diagonal pools).

**Figure 3.**
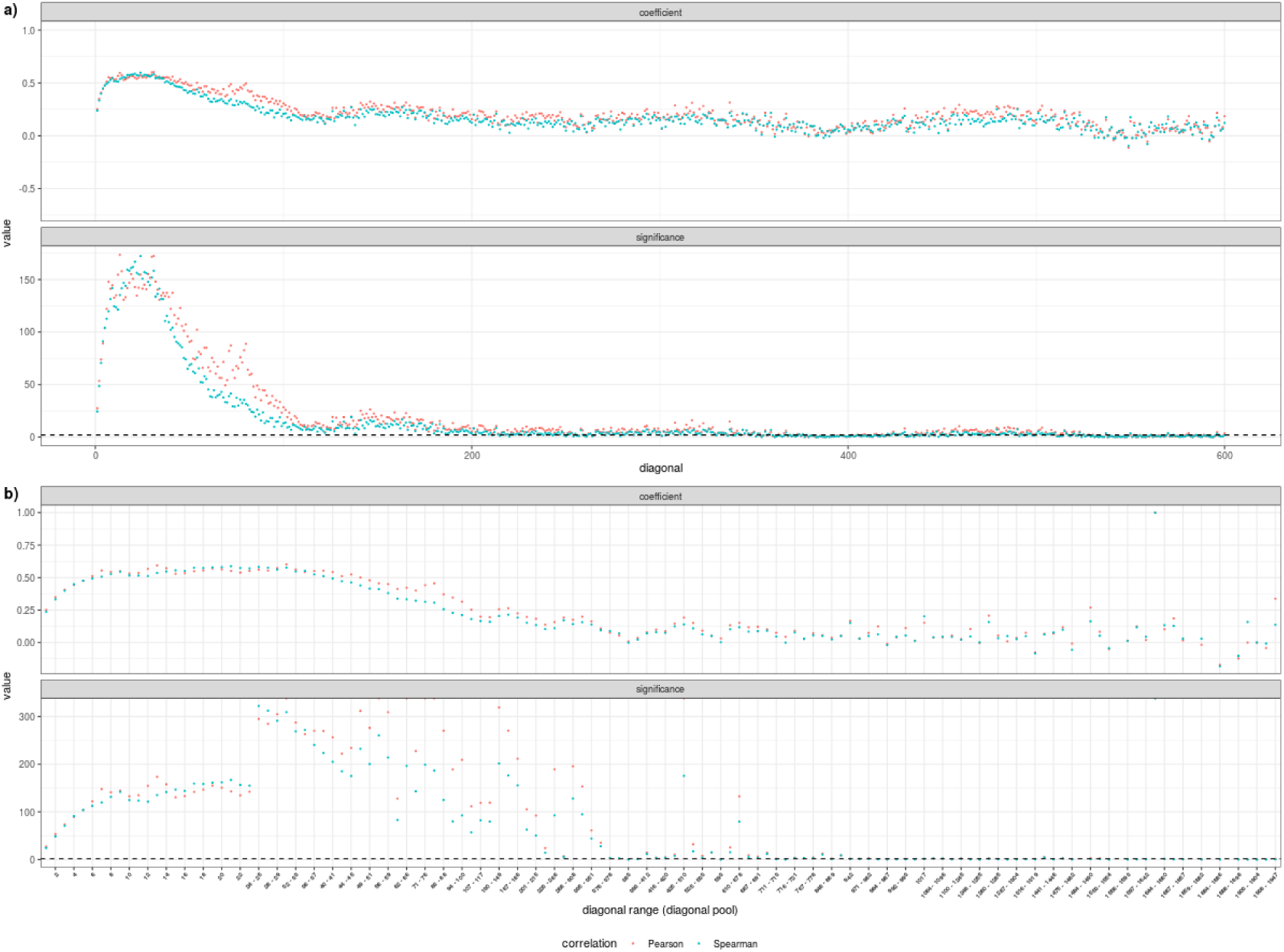
Correlations between corresponding diagonals and diagonal pools of human IMR90 and MSC cell, raw Hi-C data, chromosome 18. Pearson and Spearman correlations of paired interactions as a function of diagonal (**a**) or diagonal pool (**b**). Top panels illustrate correlation coefficient while bottom panels show respective significance expressed as negative logarithm of p-value.

We realize that maximum separation distance suggested by us comprise minor fraction of Hi-C decay range however we argue that more distant contacts are severly affected by noise and will be difficult to quantify for differential interactions anyway. This is also reflected by the fact that Hi-C data is known to follow power law distribution and the cumulative number of contacts at as little as 10 percent of diagonals comprise from 60 to even 90 percent of all contacts within the entire chromosome (Supplemental Table 2) [Lieberman-Aiden et al., 2009].

### Modelling interaction dependency using GLM

As described in previous section we observe statistically significant linear relationship between interaction intensity of two contact maps. However as can be seen in Figure 4a and 4b, small fraction of observations disobey linear pattern. We hypothesise such pairs of regions to be potential differential interactions. Due to apparent heteroscedasticity we can’t use ordinary least squares regression to model the dependency. Instead we decided to use Negative Binomial regression (see Materials and methods), which is known to perform well in analysis of overdispersed count data in general and in NGS data in particular [Hilbe, 2011, Robinson et al., 2010, McCarthy et al., 2012, Love et al., 2014]. Our model aims to capture the relationship between the number of contacts in contact map A (predictor) and number of contacts in contact map B (response) for every pair of regions given their genomic separation. Importantly, as we expect a fraction of observations to be outliers (differential interactions), we apply robust procedures in order to estimate model parameters (see Materials and methods).

**Figure 4.**
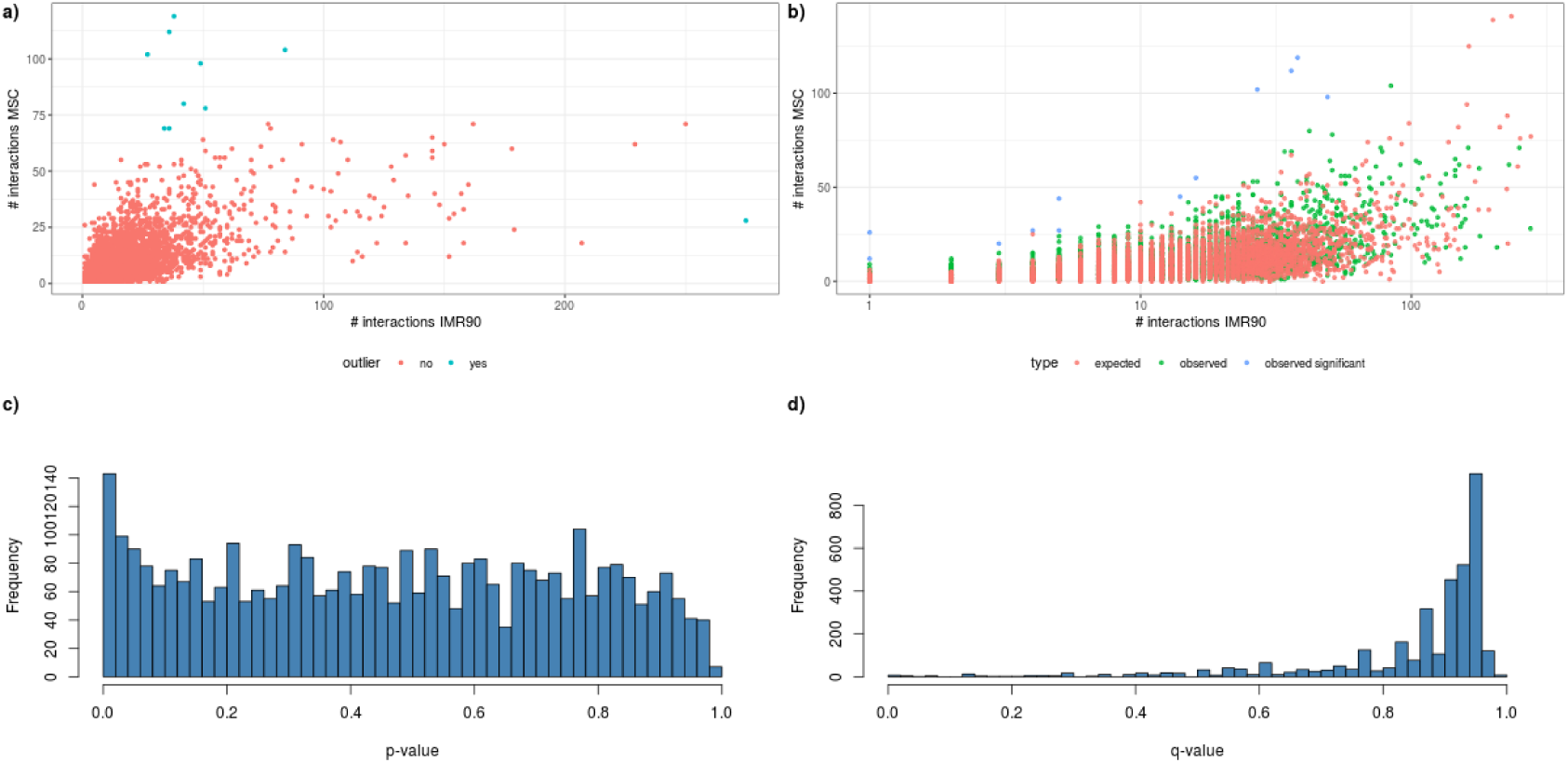
Model construction procedure. Steps a-d or b-d are performed for every diagonal from prespecified range depending on selected method of estimation. **a)** Robust regression on paired interactions set allow to determine outliers (potential differential interactions) and exclude them. **b)** The prefered method is to estimate Negative Binomial regression parameters with robust procedure rather then application of robust regression and subsequent ordinary MLE. Resulting model is used as background to calculate enrichment p-value for each paired interaction. **c)** Enrichment p-value distribution indicate deviation from null model. **d)** Finally p-values are adjusted using Benjamini-Hochberg method. Resulting q-values are used to draw significance maps.

After estimating model parameters we calculate the probability of contact enrichment of B with respect to A for every observation based on conditional distribution of B given A derived from our model (Supplemental Figure 11-16). Finally p-values are adjusted for multiple hypothesis testing using Benjamini-Hochberg method. The summary of estimation procedure is illustrated in Figure 4. The whole process is repeated for every diagonal pool from prespecified range. Results can be visualised using significance matrix where each cell express how likely given pair of regions is enriched in interactions in another Hi-C map. As there are 2 ways of choosing predictor and response the upper and lower triangles of the significance matrix correponds to each of them respectively (Figure 5c).

**Figure 5.**
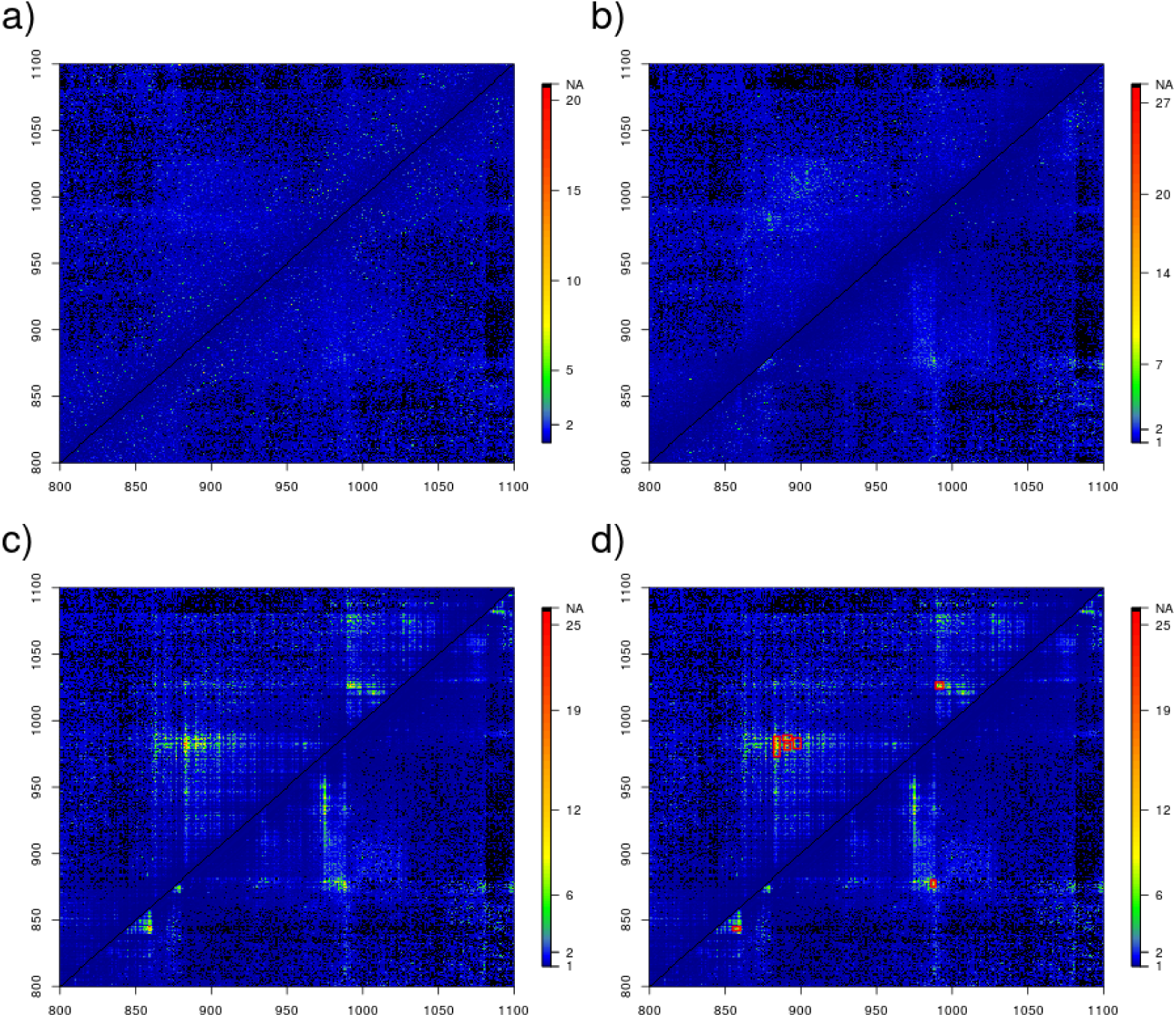
Significance maps. Every cell contain negative logarithm of adjusted p-value (i.e. q-value). Enrichment of second condition with respect to first is shown on upper triangle and the opposite on bottom triangle. **a)** IMR90 bootstrapped data 1 vs 2. **b)** IMR90 biological replicates 1 vs 2. **c)** IMR90 vs MSC. **d)** IMR90 vs MSC with LRDI regions of size at least 10.

As there is no golden standard in detection of Hi-C differential interactions and thus there is no dataset with “true” differential chromatin contacts we decided to check the performance of our method on a pair of IMR90 bootstrap and IMR90 biological replicate maps. Both of this datasets are expected to exhibit much less interaction difference as compared with different cell lines data. The results are presented in figure 5a-c and supplementary figures 17-20. The comparison of p-value distribution as well as visual inspection of significance maps provides evidence of our model’s validity.

### Detection of long range differentially interacting regions

As our testing is done on semi-diagonal (or their aggregates) separately we further reasoned that to increase the confidence of differential analysis it is straight forward to look for clusters of significant differentially interacting cells rather than individual isolated ones. Such analysis seems to be more robust against noisy data as large group of significant neighboring cells is more likely to reflect biologically relevant differential interaction as opposed to individual cell, especially for small bin sizes (high resolution). For that purpose we propose a 2 step procedure. During first step, the q-value threshold is determined, so interactions can be labelled as either significant or not. A simple method is to let the researcher specify significance threshold. Alternative approach would be an adaptive method, which uses bilinear model fit. Fitting bilinear model is based on the idea that most interactions are insignificant and therefore their negative log q-values should be small and almost constant. On the other hand we might expect a fraction of significantly enriched interactions, with their negative log q-values following a steep linear trend (see Supplemental Figure 21). In the second step significant cells are used to construct incidence matrix, which is then scanned for connected components, i.e. groups of neighboring significantly enriched cells (see Materials and methods). When algorithm completes the connected components search, a list containing smallest rectangle-like regions including significantly enriched interactions (LRDI) is returned. Resulting list may be filtered to keep only the largest LRDIs (see Figure 5d).

To asses the performance of LRDI detection method we compared the LRDI size distribution of different cell lines dataset to that from pairs of biological replicates and pairs of bootstrapped datasets. The results (Figure 6) indicate that not only the number of LRDI is higher for the dataset with different cell lines, but also detected LRDIs correspond much larger regions. Comparison of LRDI size distribution between single cell replicates versus different cells may also serve as a guidance to establish proper LRDI size cutoff for selection of biologically relevant LRDI regions.

**Figure 6.**
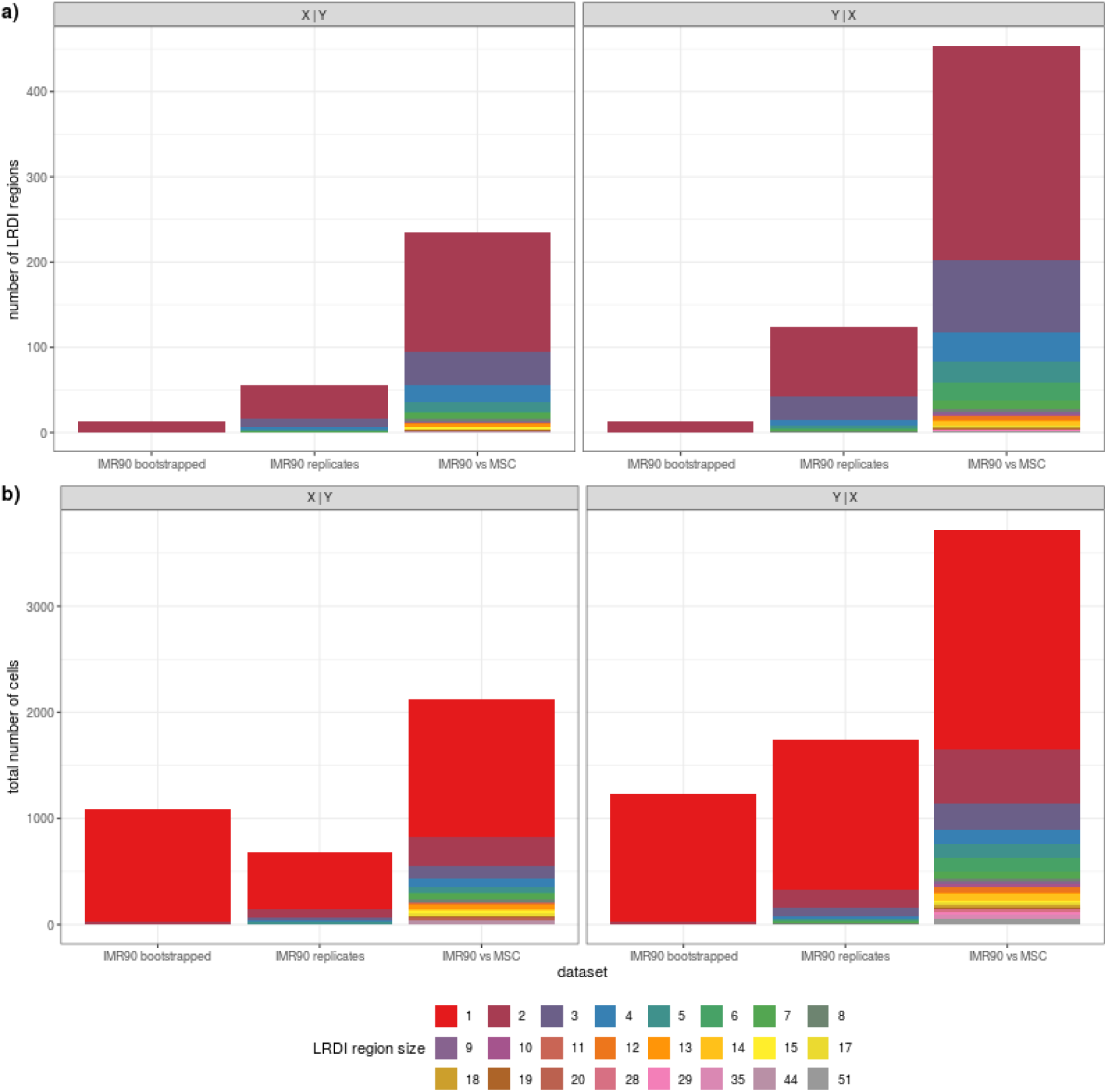
Comparison of LRDI regions size distribution. **a)** Each bar represents the number of LRDI regions of given size (i.e. including the number of significantly enriched cells) in 3 pairs of datasets: IMR90 bootstrap, IMR90 biological replicates and IMR90 vs MSC. LRDI regions of size 1 were removed for better visibility. **b)** Similar as in a), but now y axis indicates the number of LRDI regions of given size times the size of that LRDI region.

### Comparison with existing approaches

Finally we compared our method with multiple existing approaches using simulated datasets. In our benchmark we included state of the art methods designed for Hi-C differential interactions discovery. This included GLM based methods diffHiC [Lun and Smyth, 2015] and multiHiCcompare [Stansfield et al., 2019], a decay normalization based HiCcompare [Stansfield et al., 2018] and recently published SELFISH [Ardakany et al., 2019], which uses self-similarity measure. While diffHiC, multiHiCcompare and DiADeM all models the counts at bin pairs using negative binomial distribution they differ in estimation process considerably. DiffHiC requires replication for each experimental group to estimate its respective model parameters and then uses quasi likelihood F-test to determine if the group-specific models are significantly different. MultiHiCcompare acts similarily although it uses additional predictor - genomic distance between interacting bins. DiADeM only requires one replicate per group and assumes the interaction landscape is largely intact between Hi-C experiments. Model parameters are estimated from relationship of respective between group interaction abundances in pooled diagonal-wise fashion using robust procedure, which prevents the bias from entering estimates. Moreover diffHiC and multiHiCcompare fit their models using normalized data while DiADeM works with raw counts. Notably the remaining 2 methods - HiCcompare and SELFISH doesn’t require replication similarily to DiADeM. It’s worth emphasizing that we were unable to include another state of the art method FIND in this comparison due to prohibitive run times [Djekidel et al., 2018]. Nevertheless we used the approach for simulating artificial Hi-C data developed therein (see Materials and methods). However after inspection of contact decays in FIND simulated data we decided to generate additional datasets with a method developed by us, which yielded Hi-C matrices having contact decays more reminescent of real Hi-C maps (Supplemental Figure 22 and Materials and methods).

We simulated contact maps with randomly inserted differential interactions having fold change values of 2, 4, 6 and dispersion 0.1, 0.25, 0.32 (FIND simulated data) as well as real data dispersion (our method) - see Materials and methods for details. In every case we produced 2 replicates per group. For methods, which are not using replication, we pooled replicates. Finally we compared performance of every method for each fold change and dispersion value using precision-recall curves. We decided to use this metric instead of ROC curve, because the datasets are highly imbalanced, i.e. the ratio of true positive differential contacts to all possible contacts equals around 0.022 (FIND simulated data) or 0.001 (our method). The results are illustrated in figure 7. First we conclude that when fold change value is low all methods perform poorly although multiHiCcompare performs best in case of FIND simulated data and SELFISH for real data. For larger values of fold change DiADeM and multiHiCcompare seems to outperform remaining methods in case of FIND simulated data. However DiADeMs behaviour is more consistent as it performs better for real data cases. In summary we consider our approach as a good alternative for existing methods especially in absence of replication.

**Figure 7.**
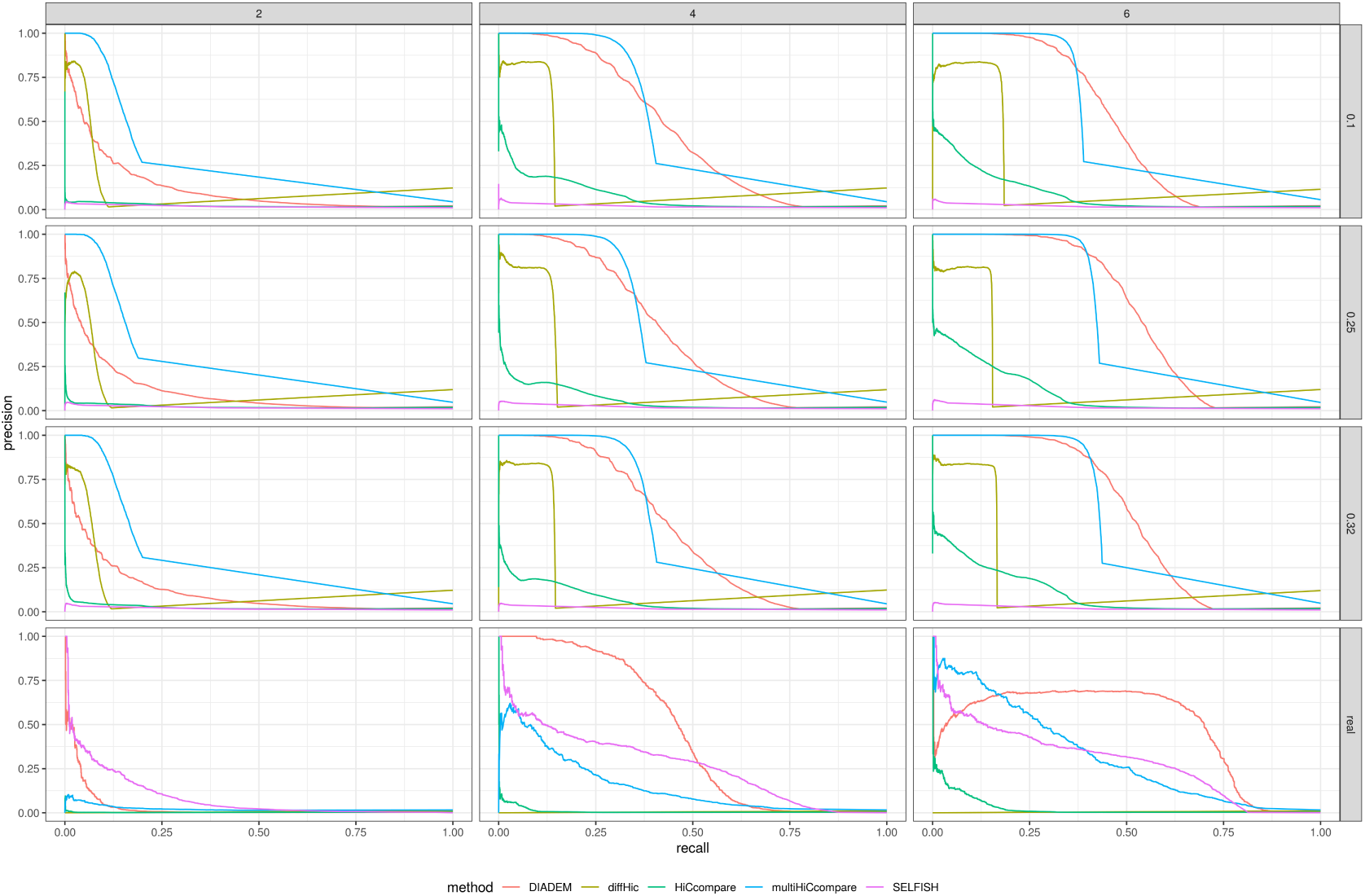
Precision-Recall curves for method evaluation on simulated data. Each column correspond to different fold change value of differential interactions while row label indicate overdispersion of simulated Hi-C datasets.

## Discussion and Conclusions

In this paper we have shown several analyses indicating that currently employed normalization techniques can affect the results of comparative analyses of multiple Hi-C datasets. What is even more worrying, the results of such analyses can be biased systematically and the direction of the bias may depend on the particular normalization technique used, giving researchers opposite results with respect to the number of enriched interactions. While this may sometimes be a minor issue (e.g. when researchers do not analyze data for differential contacts), there are situations in which there is no clear answer to the question of which normalization method is giving better results.

The natural choice in this situation is to go back to the raw, non-normalized data and see if we can test the hypothesis of interest while avoiding normalization. As we have shown, in typical cases we see a very good correlation between measurements of semi-diagonals in datasets before normalization that deteriorates after applying standard coverage uniformization. Even though this signal is only visible in the range of relatively close interactions, it happens to be the range where we are most interested in accurate prediction of differential contacts. This suggests, that the approach that might be productive would be to attempt to compare data without the uniform coverage normalization.

Since we still need to have some sort of mapping between the raw counts that would take into account the global effects such as the total read count and the observable variance differences, we chose to use the generalized linear models approach to represent the relationship between counts found in the compared Hi-C experiments for each semi-diagonal of the compared maps. This allowed us to have a relatively flexible model, that enables for representation of the heteroscedacticity of the data and the expected contact decay with increased distance while avoiding the deterioration of the signals stemming from over-normalization of the coverages.

Using this model, we are able to not only identify regions that are significantly different in their contact frequency, but also aggregate them into larger regions that are likely to coincide with biologically relevant events. We show that the proposed approach gives much more differential contacts in comparison of biologically different samples that betwen replicates and that the detected differential regions are also significantly larger. We have implemented this approach in the software tool called DIADEM and made it available to the public.

In summary, we consider the results very promising. We hope that the fact that we documented the significant side-effects of the widely used normalization techniques will lead to better understanding of the results of Hi-C analyses. Even more importantly, we hope that the solution to the identified problem that we provide will lead to wide adoption of the methods we propose and in turn to more unbiased results in future Hi-C analyses.

## Materials and methods

### Software availability

The software is freely available as R package DIADEM under MIT license at: https://github.com/rz6/DIADEM.

#### Data

The Hi-C data used in this study comes from publicly available GEO repositories, studies: GSE63525, GSE52457, GSE87112 and GSE96107.

#### Definitions

Throughout the article we use following notation:

- Hi-C contact map *X*:

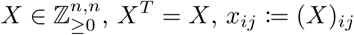
- *k*^th^ diagonal elements of contact map *X*:

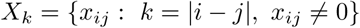
- paired interaction set 𝒟_*k*_ of matrices *X, Y* at genomic distance *k*:

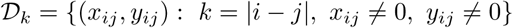 The set 𝒟_*k*_ can be considered as a realization of (bivariate) probability distribution *F*_*k*_. We also use the notation **x**^(*k*)^, **y**^(*k*)^ in order to refer to *X, Y* dataset elements of 𝒟_*k*_ respectively, i.e.: 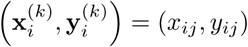.

### Coverage calculation

The coverage of contact map *X* is defined as a function mapping a bin with the sum of its contacts:

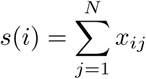

### Contact decay calculation

The contact decay of contact map *X* is defined as a function mapping diagonal with the mean of its contacts:

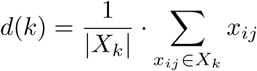

### Diagonal correlations calculation

The Pearson *r*, Spearman *ρ* and Kendall *τ* correlation coefficients are calculated between **x**^(*k*)^ and **y**^(*k*)^ vectors for varying values of *k*.

### Diagonal pooling

Decay pooling process is based on testing following null hypothesis:

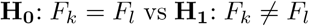

In order to assert the above hypothesis we employ distribution free, approximate permutation test developed by [Székely et al., 2004]. The test is based on E-statistic (energy distance) and is implemented in R package energy [Rizzo and Szekely, 2019]. The E-statistic measures the differences in sum of interpoint distances between and within 2 multivariate samples 𝒟_*k*_, 𝒟_*l*_ of sizes *n, m* respectively:

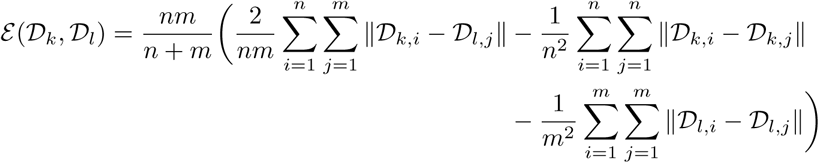

Large values of *ε* indicate strong evidence against null hypothesis. The approximate null distribution of *ε* is obtained by bootstrapping pooled sample *W* = 𝒟_*k*_ ∪ 𝒟_*l*_ (s.t. |*W*| = *n* + *m*). More precisely *B* bootstrap samples *W* ^(1)^, *W* ^(2)^, …, *W* ^(*B*)^ are generated by randomly permuting *W*. Afterwards each sample *W*^*b*^ is divided on two subsamples 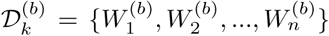 and 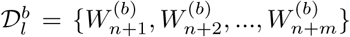 and the corresponding 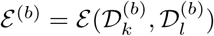 is calculated. Finally the p-value is calculated according to:

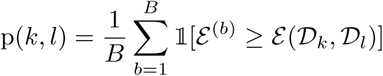

Now the diagonal pool *p* consists of paired interaction sets corresponding to consecutive diagonals 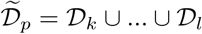 such that at given significance level *α*:

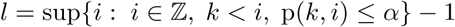

In other words given 𝒟_*k*_ we seek for first diagonal *i* such that we can reject the null hypothesis: *F*_*k*_ = *F*_*i*_. The paired interaction sets 𝒟_*k*_ through 𝒟_*l*_ such that *l* = *i* − 1 comprise pool *p* and 𝒟_*i*_ enters next pool, i.e. *p* + 1, which range is to be determined analogously.

The reason for this choice of pooling arise from the existance of contact decay - lower the genomic distance between interacting regions, higher the mean number of contacts. Correspondingly if *i < j < k* one might expect the following relation *ε* (𝒟_*i*_, 𝒟_*k*_) ≥ *ε* (𝒟_*j*_, 𝒟_*k*_) to be satisfied for majority of cases.

### Model definition and estimation

Given pooled, paired interaction set 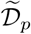, which corresponds to experimental conditions *X* and *Y* we assume the number of contacts 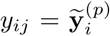 can be predicted by the abundance of 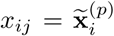 using generalized linear model with negative binomial distribution *Y* |*X* ∼ *NB*(*µ*_*i*_, *ϕ*_*p*_) given no differential interactions exist between the two datasets:

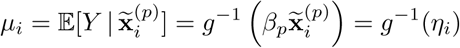

Accordingly model parameters to be estimated are expressed as: *θ*_*p*∈{1,…,*P*}_ = (*β*_*p*_, *ϕ*_*p*_) where *P* is the number of diagonal pools. However parameter estimation is not straight forward, because a fraction of observations (alleged differential interactions) are expected to follow from contamination process, which is unlikely to be distributed as the “no-difference” model. Although these outlying observations are rare they may still bias the parameter estimates if not accounted for properly.

Below we present 2 possible solutions intended to provide robustness against outliers during parameter estimation. In our opinion the first method should be prefered as it was designed for particular case of robust estimation of NB GLM parameters, which is the exact case in our problem. However the approach is computationally demanding and in some cases, especially when analyzing high resolution, large Hi-C datasets may turn out to be prohibitive. For that situation we suggest alternative solution based on outlier detection and removal followed by ordinary MLE.

#### Robust estimation

The techniques for unbiased parameter estimation in presence of outliers or small departures from model parameters are termed as robust. In this work we use the robust M-estimators for NB GLM parameters derived by [Aeberhard et al., 2014]. More precisely robust estimators of *β*_*p*_ and *ϕ*_*p*_ can be obtained by solving the following estimating equations:

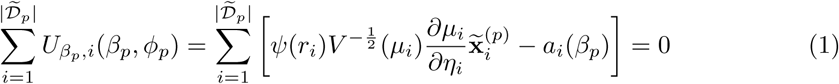

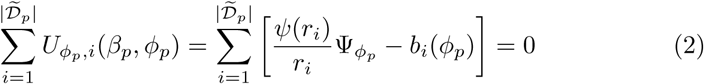

where:

- 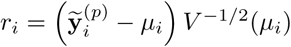 is a Pearson residual,
- 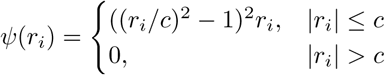 is a Tukey’s biweight function with tuning constant *c*, which express the tradeoff between robustness and efficiency. In our method *c* = 4 as suggested in [Aeberhard et al., 2014] paper,
- 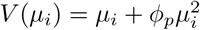 is the conditional variance,
- 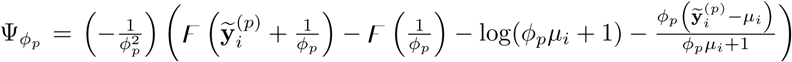is a derivative of the log likelihood function with respect to *ϕ*_*p*_ and *F* (*u*) = *∂* log Γ(*u*)*/∂u* is the digamma function,
- 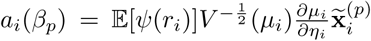 and 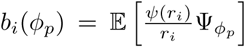 are correction terms providing Fisher consistency of estimators.

The algorithm to solve equations 1 and 2 proceeds by repeatedly finding 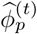 from 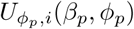 given 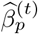 and then solving 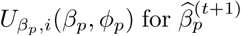 while fixing 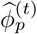. The iteration is repeated until attaining convergence in terms of 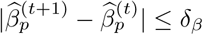 and 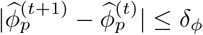. Initial estimates for 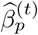 are obtained by fitting Poisson GLM.

#### Removal of outliers followed by MLE

In order to determine outliers we employ robust regression approach using SMDM-estimator developed by [Koller and Stahel, 2011], which consists of following estimation steps:

1. S-estimation,
2. M-estimation,
3. D-estimation (Design Adaptive Scale Estimate),
4. M-estimation.

The combination of first two steps is known as MM-estimation and consists of robust estimation of scale (i.e. the variance of error term), which is then used in the process of estimating location (i.e. the regression coefficients). D-estimation is a novel scale estimator derived by [Koller and Stahel, 2011], which is obtained by solving following equation for *σ*_*D*_:

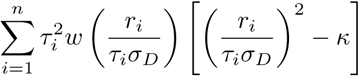

In the above equation:

- *w*(*r*) = *ψ* (*r*)*/r* is a weighting function with *ψ* - Huber psi, Tukey biweight, or different loss function,
- *κ* ensures Fisher consistency at the model,
- *r*_*i*_ is *i*-th residual,
- *τ*_*i*_ are correction factors, which are designed to reflect the heteroskedasticity of the distributions of the residuals *r*_*i*_ and depend on the leverage of *i*-th observation and the choice of *ψ* function.

As the exact distribution of residuals is unknown it is approximated using *von Mises expansion* of 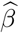 and the precise values of *r*_*i*_ are calculated based on estimates obtained in previous steps. For more details regarding the definition of *τ*_*i*_ and calculation of *r*_*i*_ one should refer to [Koller and Stahel, 2011]. Finally during last step regression coefficients are reestimated using *σ*_*D*_. The robustness weights produced during the final M-step are applied to determine outliers - a 0 weight observations. Described method is implemented in R package robustbase [Maechler et al., 2019].

### Significance calculation

Given paired interaction 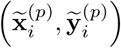 and parameter estimates for diagonal pool 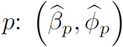we calculate the probability of observing the number of interactions *Y* between genomic loci *i* and *j* at least as abundant as 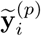 given 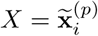 according to following formula:

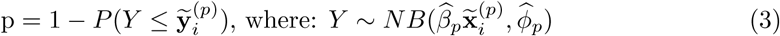

As this test is performed for multiple pairs of regions in selected chromosome we correct resulting p-values using Benjamini-Hochberg procedure and obtain q-values.

### Detection of LRDI regions

In order to determine long range differentially interacting (LRDI) regions a significance level *α* is required. The choice of *α* is made by either manually selecting arbitrary value like standard 0.05 or 0.01 thresholds or by automated procedure. The automated approach works by first arranging negative logarithm of q-values in ascending order and then uses obtained vector to fit a bilinear model. Resulting linear functions are associated with non-differential and differential interaction regimes of q-values. The intersection point between linear functions is used to determine *α* threshold.

After selecting *α* q-value of every pair of regions is compared with *α* and labelled as differential or not resulting in binary 1-0 matrix. This matrix is converted to adjacency list of a graph induced by chromosomal regions (contact map bins). The graph is scanned for connected components (LRDI regions) using standard algorithm implemented in R package raster [Hijmans, 2018]. Briefly the algorithm works by iterating through list of vertices and invoking Depth First Search procedure for every unvisited vertex until all vertices are marked as visited.

### Generation of simulated data

The artificial differential interactions data were simulated in two ways. First we used the method implemented in FIND [Djekidel et al., 2018], which produces simulated Hi-C contact maps given real Hi-C matrices, dispersion and fold changes. We used IMR90 contact maps (2 technical replicates) from [Rao et al., 2014] and MSC contact maps (2 technical replicates) from [Dixon et al., 2015] with dispersion values 0.1, 0.25, 0.32 and fold changes of 2, 4, and 6.

Second method is based on the application of our model. First we estimate two NB GLM models using the method developed by us on IMR90 versus MSC, first replicates data - model 1 and IMR90 versus MSC second replicates data - model 2. Then we generate two pairs of replicates data by:

- simulating Hi-C matrix 1 from model 1 given the first replicate of IMR90,
- simulating Hi-C matrix 2 from model 2 given the second replicate of IMR90.

To obtain the second simulated data replicates we input the model with MSC data, replicates 1 and 2 respectively. As a result we obtain 2 pairs of contact maps and refer to each pair as simulated data replicates. Afterwards we randomly select 200 cells (100 in first pair and 100 in second pair of simulated replicates) and multiply the number of interactions in selected cells as well as all cells neighboring them by specified fold change value. Moreover the process of random selection of cells is restricted by the minimum distance of selected cells and specific range of diagonals. More precisely any pair of randomly selected cells is guaranteed to be separated by at least 10 cells and all randomly selected cells are within second and 120^th^ diagonal. The described process of simulating Hi-C data ensures that obtained matrices preserve important characteristics of real Hi-C datasets like coverage and contact decay.

In order to compare the performance of methods we calculate precision recall curves using R package ROCR [Sing et al., 2005].

## Acknowledgments

This work was supported by the Polish National Science Centre grant decision No. [DEC 2015/16/W/NZ2/00314].

## References

Aeberhard, W. H., Cantoni, E., and Heritier, S. (2014). Robust inference in the negative binomial regression model with an application to falls data. Biometrics, 70(4):920–931.

Ardakany, A. R., Ay, F., and Lonardi, S. (2019). Selfish: discovery of differential chromatin interactions via a self-similarity measure. Bioinformatics, 35(14):i145–i153.

Dekker, J., Rippe, K., Dekker, M., and Kleckner, N. (2002). Capturing chromosome conformation. science, 295(5558):1306–1311.

Dixon, J. R., Jung, I., Selvaraj, S., Shen, Y., Antosiewicz-Bourget, J. E., Lee, A. Y., Ye, Z., Kim, A., Rajagopal, N., Xie, W., et al. (2015). Chromatin architecture reorganization during stem cell differentiation. Nature, 518(7539):331.

Djekidel, M. N., Chen, Y., and Zhang, M. Q. (2018). Find: differential chromatin interactions detection using a spatial poisson process. Genome research, 28(3):412–422.

Eagen, K. P. (2018). Principles of chromosome architecture revealed by hi-c. Trends in biochemical sciences, 43(6):469–478.

Hijmans, R. J. (2018). raster: Geographic Data Analysis and Modeling. R package version 2.8-4.

Hilbe, J. M. (2011). Negative binomial regression. Cambridge University Press.

Hu, M., Deng, K., Selvaraj, S., Qin, Z., Ren, B., and Liu, J. S. (2012). Hicnorm: removing biases in hi-c data via poisson regression. Bioinformatics, 28(23):3131–3133.

Imakaev, M., Fudenberg, G., McCord, R. P., Naumova, N., Goloborodko, A., Lajoie, B. R., Dekker, J., and Mirny, L. A. (2012). Iterative correction of hi-c data reveals hallmarks of chromosome organization. Nature methods, 9(10):999.

Koller, M. and Stahel, W. A. (2011). Sharpening wald-type inference in robust regression for small samples. Computational Statistics & Data Analysis, 55(8):2504–2515.

Lieberman-Aiden, E., Van Berkum, N. L., Williams, L., Imakaev, M., Ragoczy, T., Telling, A., Amit, I., Lajoie, B. R., Sabo, P. J., Dorschner, M. O., et al. (2009). Comprehensive mapping of long-range interactions reveals folding principles of the human genome. science, 326(5950):289–293.

Love, M. I., Huber, W., and Anders, S. (2014). Moderated estimation of fold change and dispersion for rna-seq data with deseq2. Genome biology, 15(12):550.

Lun, A. T. and Smyth, G. K. (2015). diffhic: a bioconductor package to detect differential genomic interactions in hi-c data. BMC bioinformatics, 16(1):258.

Maechler, M., Rousseeuw, P., Croux, C., Todorov, V., Ruckstuhl, A., Salibian-Barrera, M., Verbeke, T., Koller, M., Conceicao, E. L. T., and Anna di Palma, M. (2019). robustbase: Basic Robust Statistics. R package version 0.93-5.

McCarthy, D. J., Chen, Y., and Smyth, G. K. (2012). Differential expression analysis of multifactor rna-seq experiments with respect to biological variation. Nucleic acids research, 40(10):4288–4297.

Nagano, T., Várnai, C., Schoenfelder, S., Javierre, B.-M., Wingett, S. W., and Fraser, P. (2015). Comparison of hi-c results using in-solution versus in-nucleus ligation. Genome biology, 16(1):175.

Rada-Iglesias, A., Grosveld, F. G., and Papantonis, A. (2018). Forces driving the three-dimensional folding of eukaryotic genomes. Molecular systems biology, 14(6):e8214.

Rao, S. S., Huntley, M. H., Durand, N. C., Stamenova, E. K., Bochkov, I. D., Robinson, J. T., Sanborn, A. L., Machol, I., Omer, A. D., Lander, E. S., et al. (2014). A 3d map of the human genome at kilobase resolution reveals principles of chromatin looping. Cell, 159(7):1665–1680.

Rizzo, M. and Szekely, G. (2019). energy: E-Statistics: Multivariate Inference via the Energy of Data. R package version 1.7-6.

Robinson, M. D., McCarthy, D. J., and Smyth, G. K. (2010). edger: a bioconductor package for differential expression analysis of digital gene expression data. Bioinformatics, 26(1):139–140.

Rudan, M. V., Barrington, C., Henderson, S., Ernst, C., Odom, D. T., Tanay, A., and Hadjur, S. (2015). Comparative hi-c reveals that ctcf underlies evolution of chromosomal domain architecture. Cell reports, 10(8):1297–1309.

Schmitt, A. D., Hu, M., Jung, I., Xu, Z., Qiu, Y., Tan, C. L., Li, Y., Lin, S., Lin, Y., Barr, C. L., et al. (2016). A compendium of chromatin contact maps reveals spatially active regions in the human genome. Cell reports, 17(8):2042–2059.

Sing, T., Sander, O., Beerenwinkel, N., and Lengauer, T. (2005). Rocr: visualizing classifier performance in r. Bioinformatics, 21(20):7881.

Stansfield, J. C., Cresswell, K. G., and Dozmorov, M. G. (2019). multihiccompare: joint normalization and comparative analysis of complex hi-c experiments. Bioinformatics.

Stansfield, J. C., Cresswell, K. G., Vladimirov, V. I., and Dozmorov, M. G. (2018). Hiccompare: an r-package for joint normalization and comparison of hi-c datasets. BMC bioinformatics, 19(1):279.

Székely, G. J., Rizzo, M. L., et al. (2004). Testing for equal distributions in high dimension. InterStat, 5(16.10):1249–1272.

Venables, W. N. and Ripley, B. D. (2002). Modern Applied Statistics with S. Springer, New York, fourth edition. ISBN 0-387-95457-0.

Yaffe, E. and Tanay, A. (2011). Probabilistic modeling of hi-c contact maps eliminates systematic biases to characterize global chromosomal architecture. Nature genetics, 43(11):1059.

