## Supplementary data for "DiADeM: differential analysis via dependency modelling of chromatin interactions with robust generalized linear models" (from another version of this preprint: version 2, 2019-12-16)

### Supplementary Information for the paper entitled ”DIADEM: differential analysis via dependency modelling of chromatin interactions with robust generalized linear models”

Rafal Zaborowski    Bartek Wilczynski

December 12, 2019

### 1 Supplementary figures

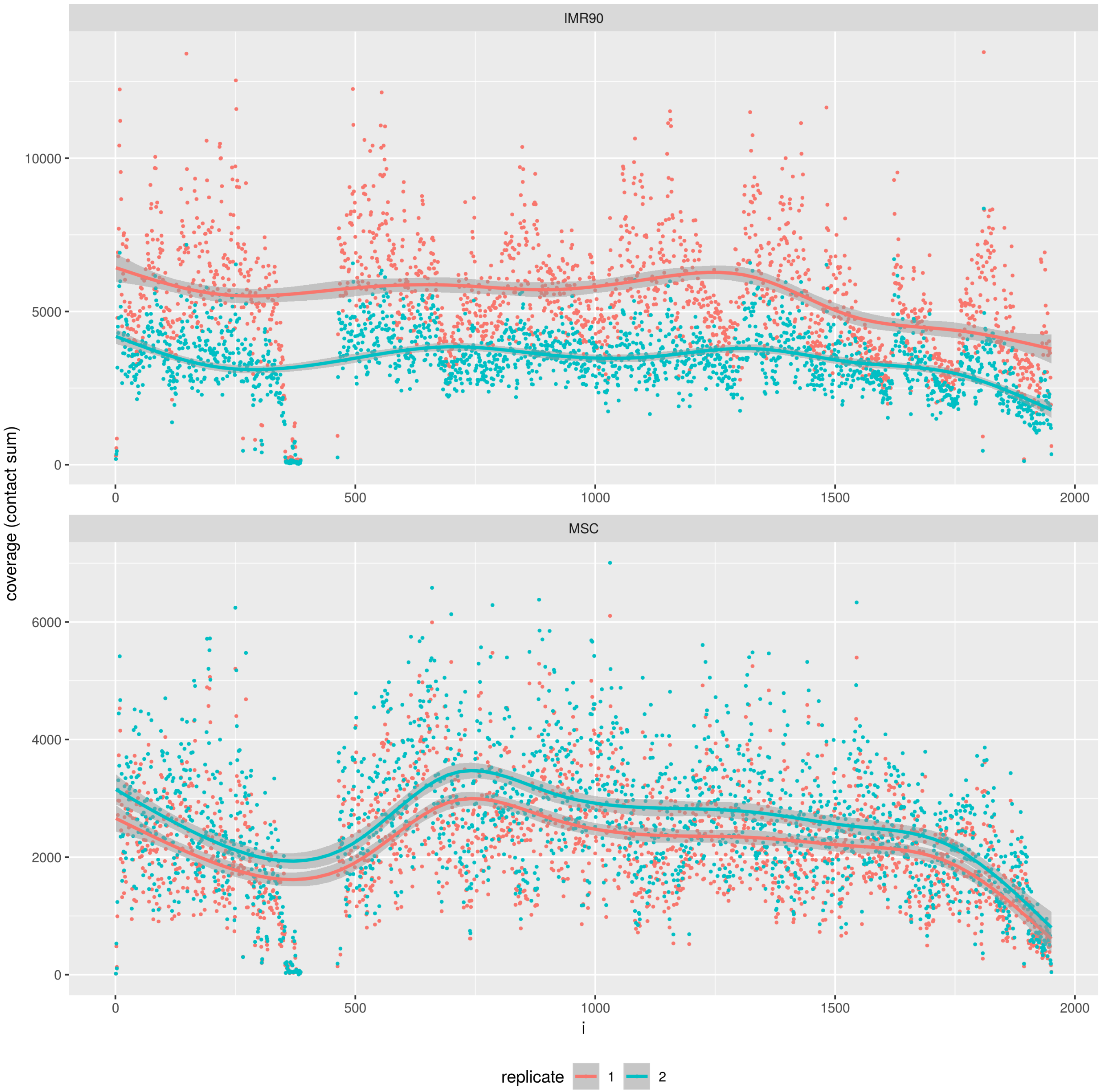


Figure 1: **Coverage comparison between biological replicates of human Hi-C data from IMR90 and MSC cell lines, chromosome 18.**

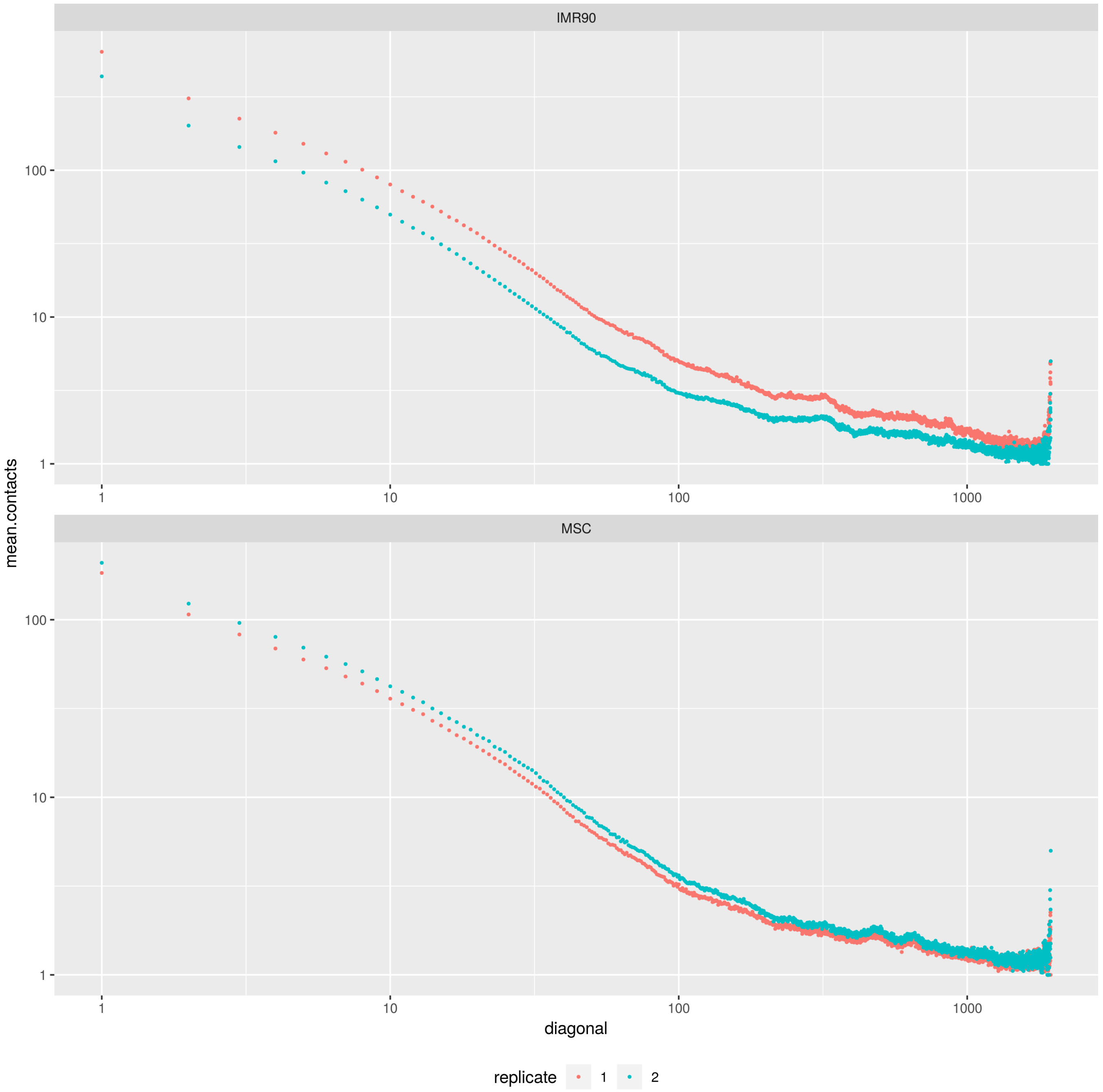


Figure 2: **Contact decay comparison between biological replicates of human Hi-C data from IMR90 and MSC cell lines, chromosome 18.**

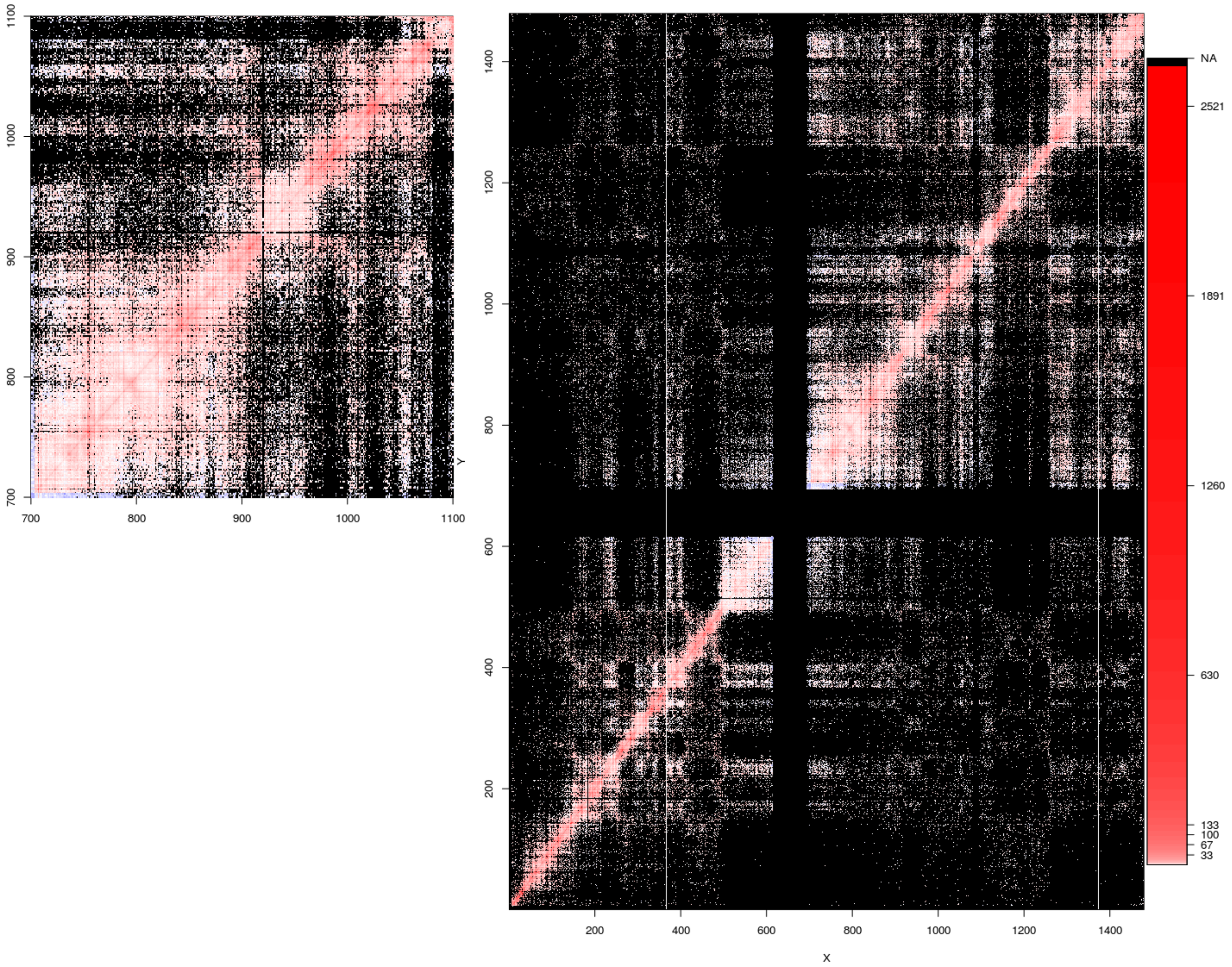


Figure 3: **Fold change map of human raw Hi-C data IMR90 / MSC cell lines, chromosome 19.**

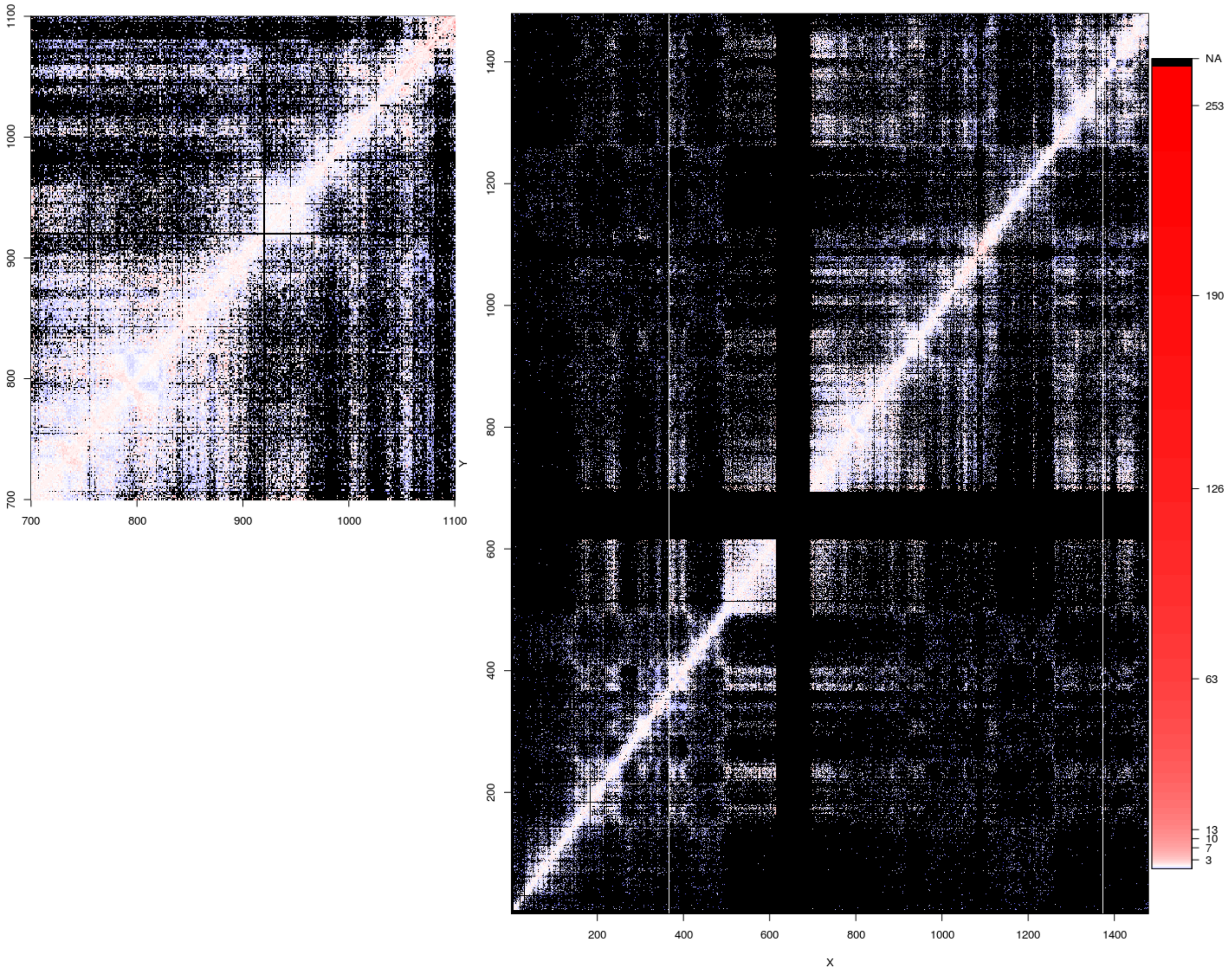


Figure 4: **Fold change map of human ICE-corrected Hi-C data IMR90 / MSC cell lines, chromosome 19.**

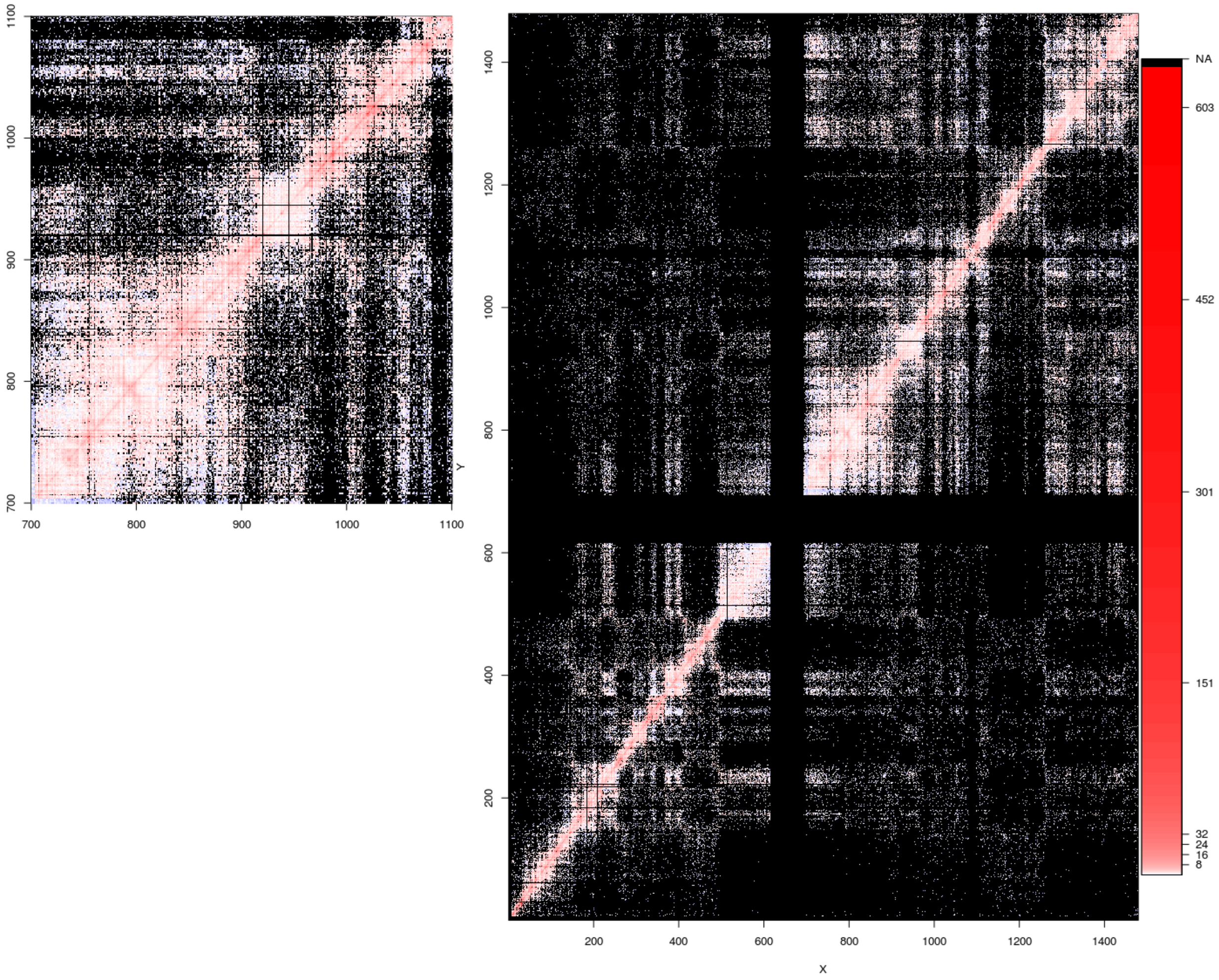


Figure 5: **Fold change map of human HiCnorm-corrected Hi-C data IMR90 / MSC cell lines, chromosome 19.**

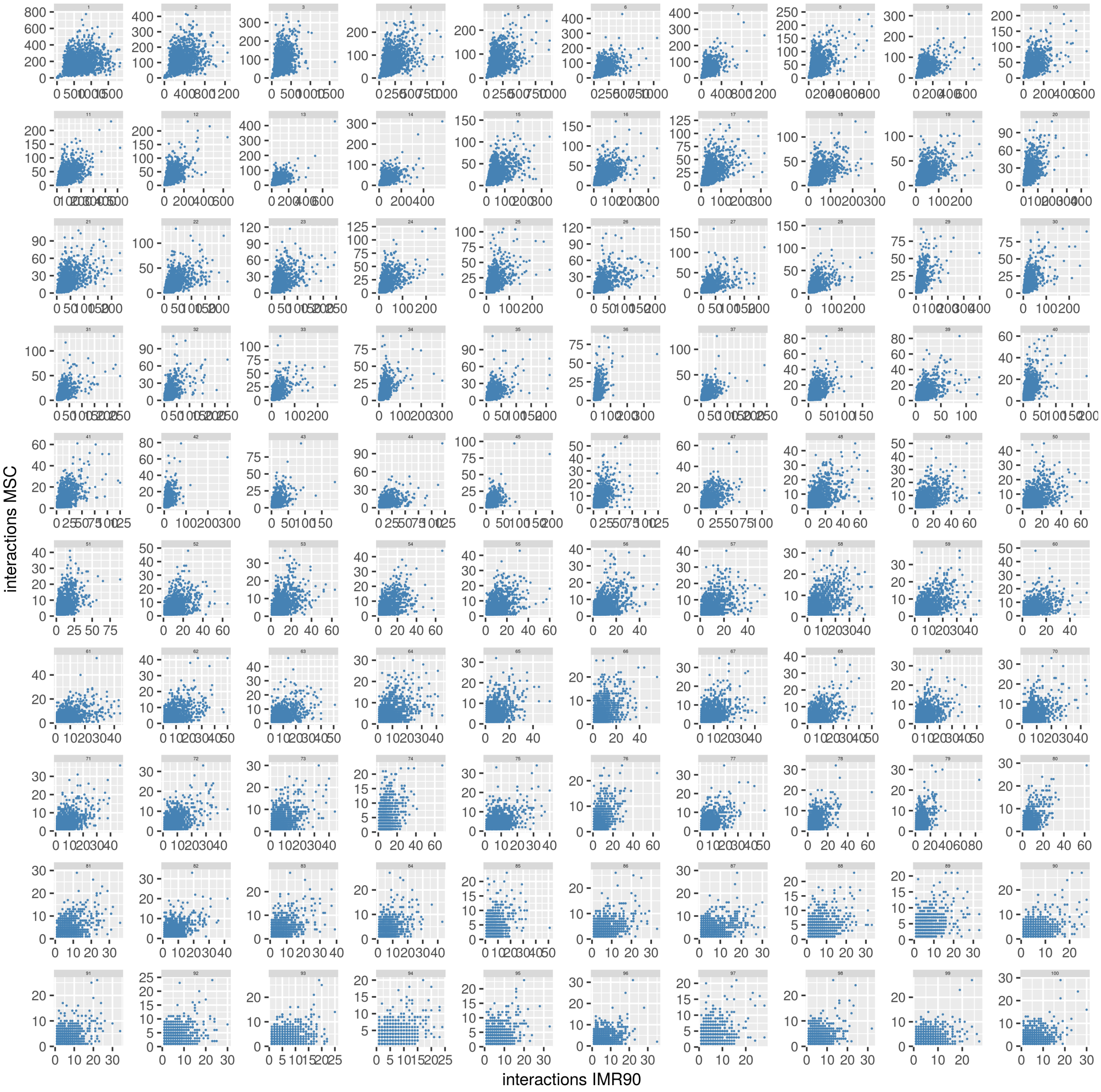


Figure 6: **Dependency between the number of Hi-C interactions in IMR90 and MSC cell lines.** Every point (paired interaction) is associated with a pair of genomic regions in human chromosome 18. It represents the relationship between the number of interactions in IMR90 dataset (X axis) vs MSC dataset (Y axis) at given pair of loci. Paired interactions are categorized by diagonals, i.e. the linear genomic distance between interacting regions.

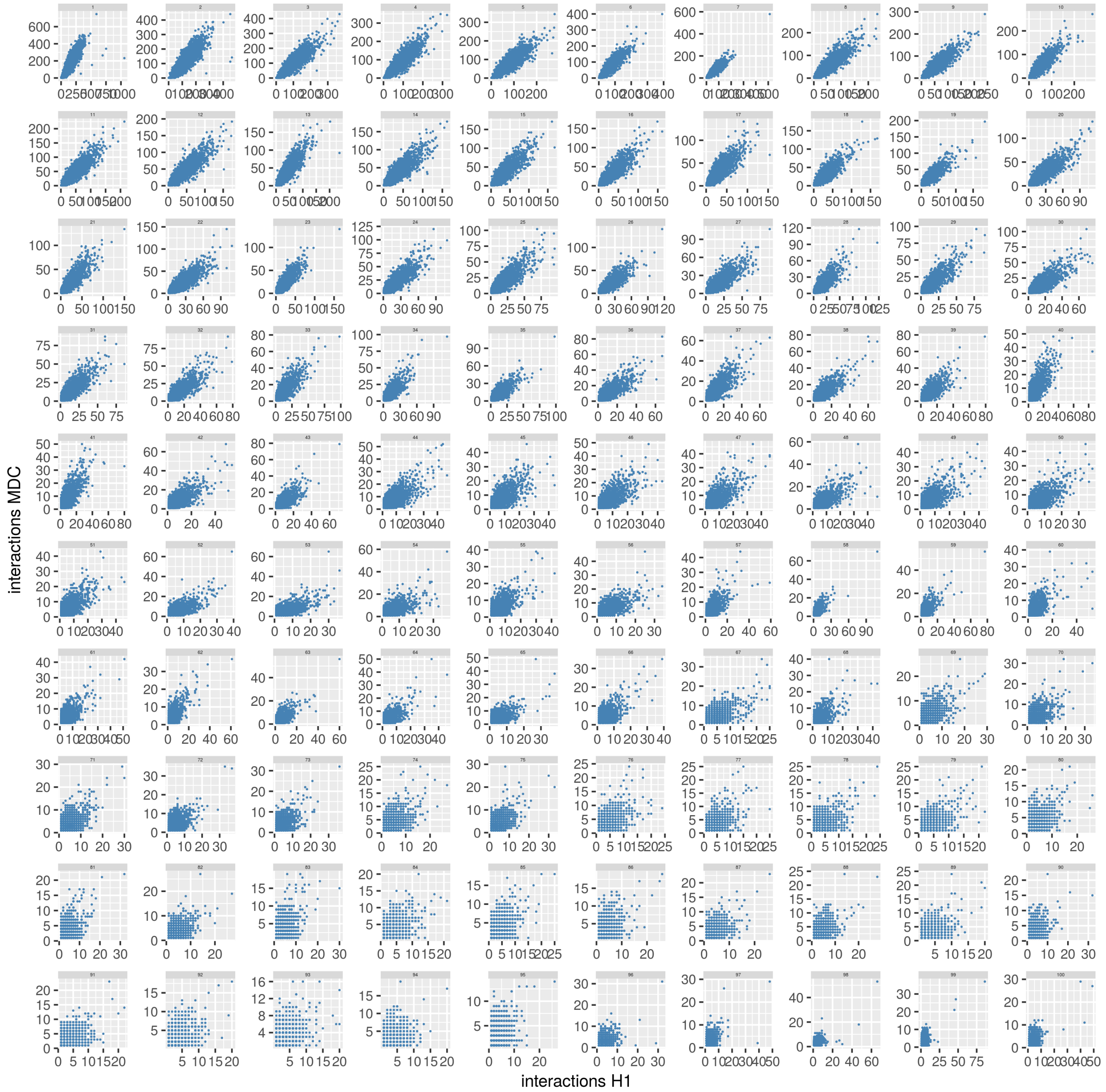


Figure 7: **Dependency between the number of Hi-C interactions in H1 and MDC cell lines.** The same as in fig. 6, but the datasets are H1 vs MDC cell lines, chromosome 10.

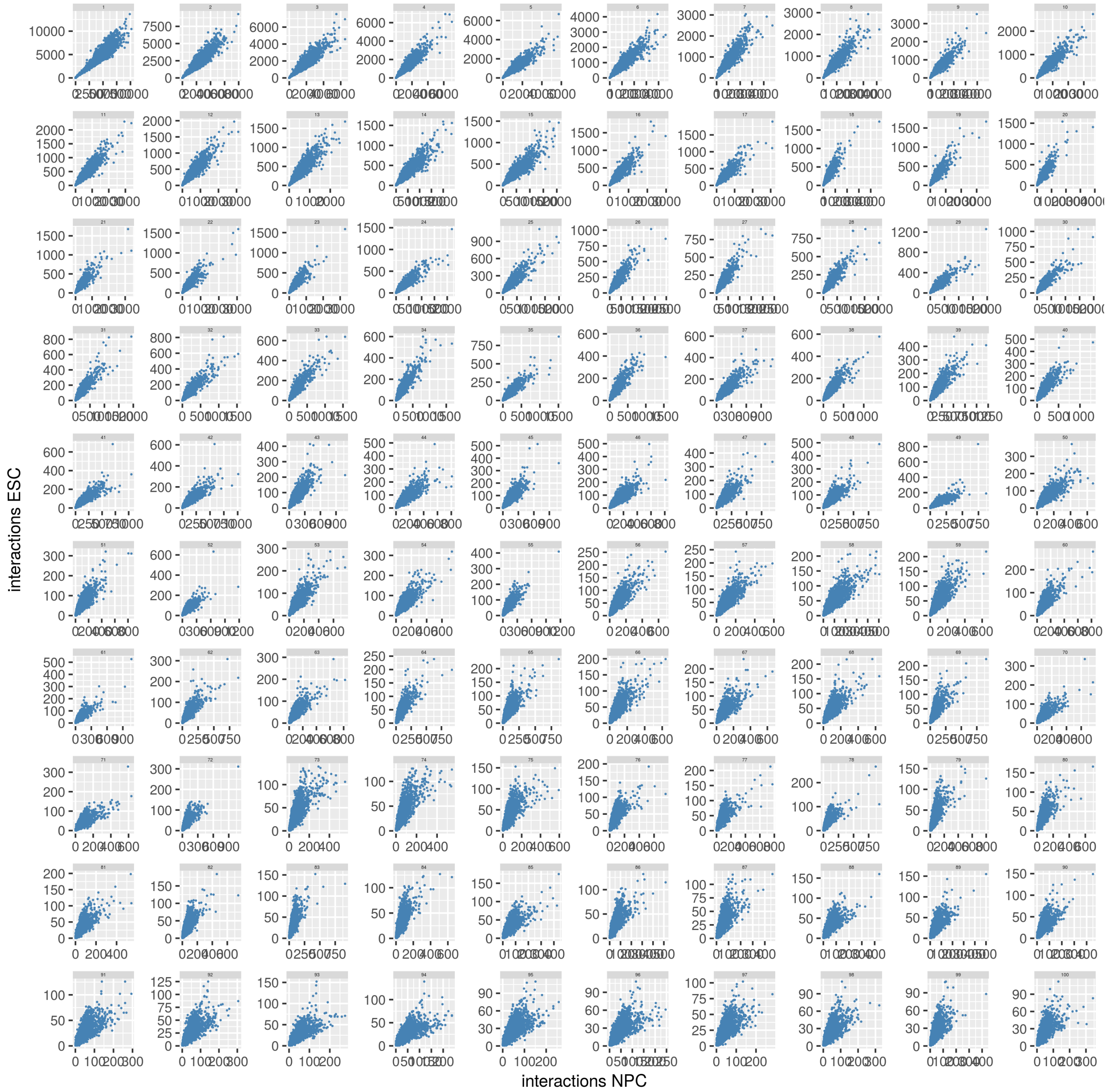


Figure 8: **Dependency between the number of Hi-C interactions in NPC and ESC cell lines.** The same as in fig. 6, but the datasets are NPC vs ESC mouse cell lines, chromosome 4.

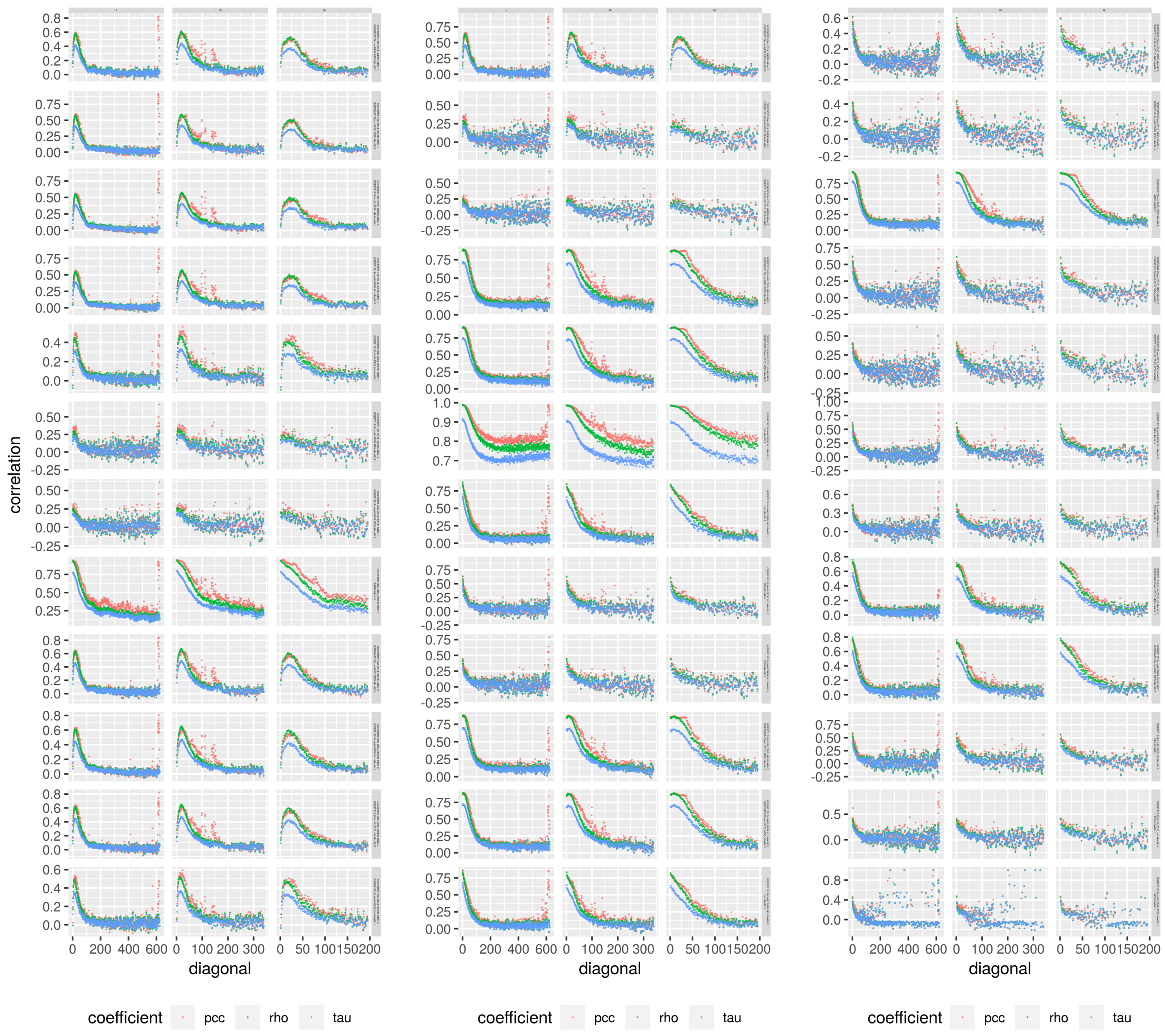


Figure 9: **Between datasets diagonal-wise correlations.** Every point express the correlation between interactions from a pair of Hi-C datasets (contact maps) at given separation between interacting loci refered to as diagonal. Only first 10% of selected chromosomes diagonals are shown. Remaining diagonal correlations are insignificant. Shown is the data for chromosomes 1,10,18 from 36 pairs of Hi-C datasets. 3 correlation measures were calculated: Pearson, Spearman and Kendall.

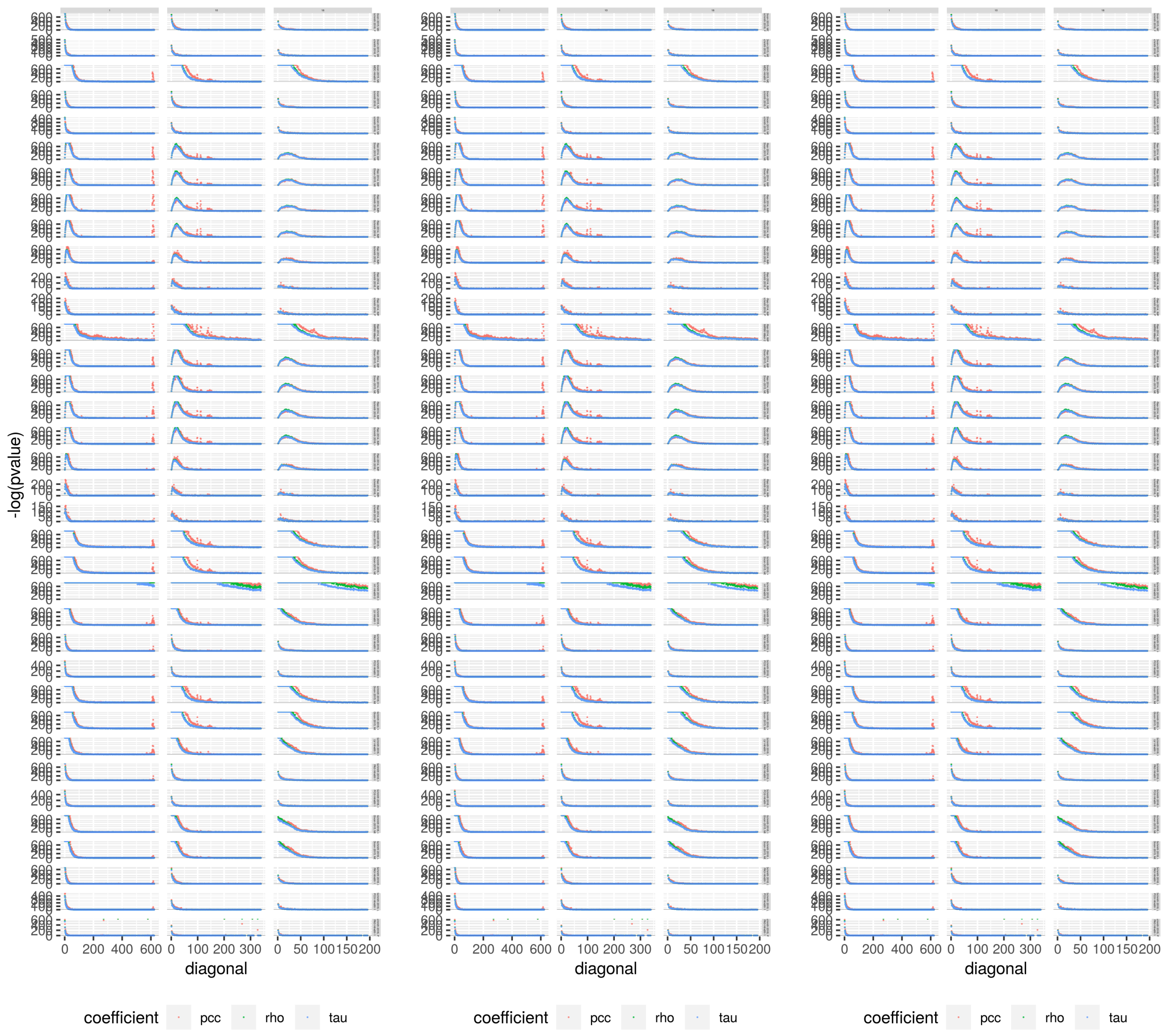


Figure 10: **Between datasets diagonal-wise correlation significances.** The same as in fig. 9, but significances of corresponding correlations are shown.

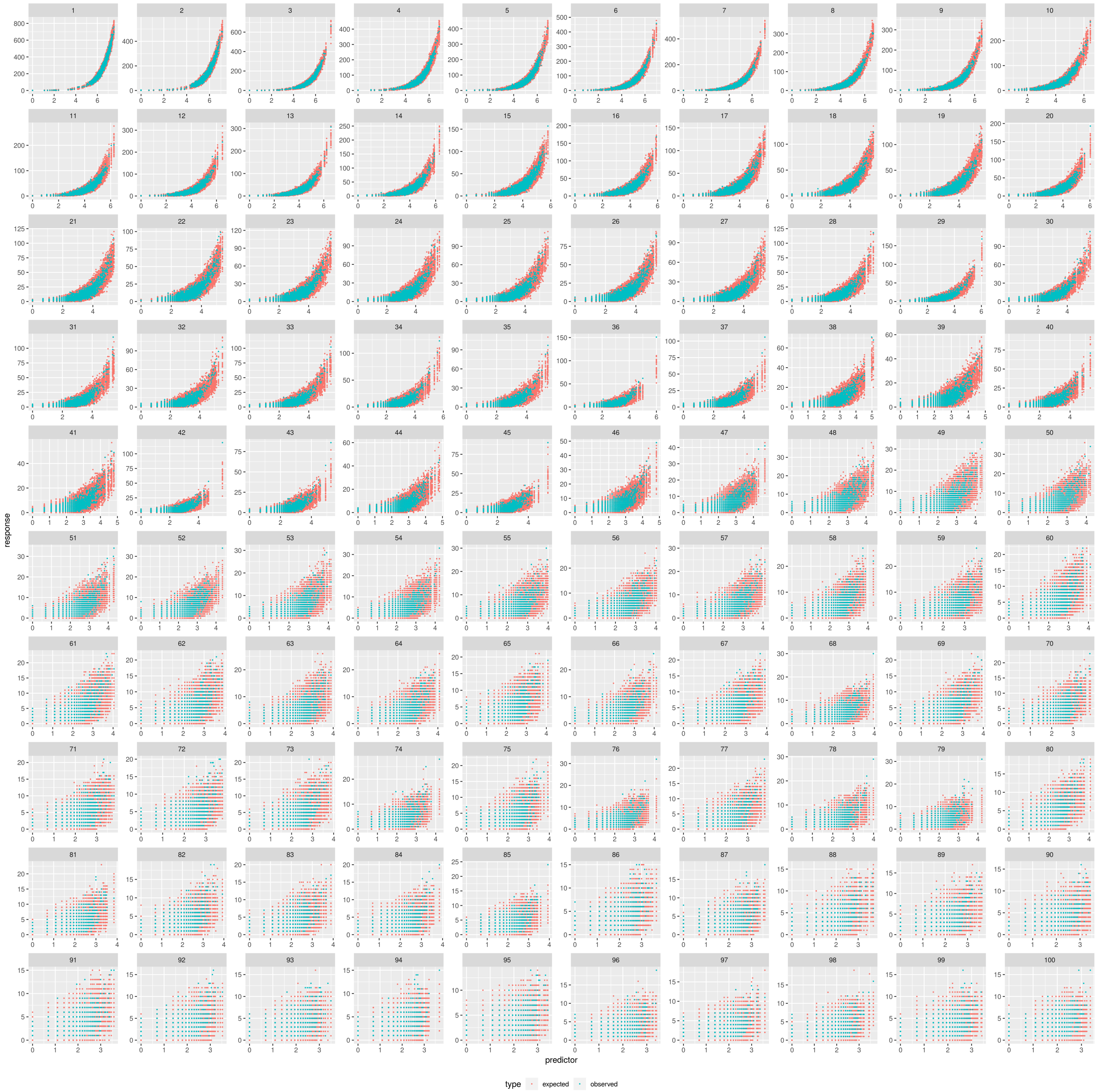


Figure 11: **Expected vs observed paired interactions for IMR90 bootstrapped data.** Each point (i.e. paired interaction) is associated with a pair of interacting regions in a Hi-C contact map with point coordinates being: the number of interactions in first contact map (X coordinate) and the number of interactions in second contact map (Y coordinate). Paired interactions are categorized by diagonal. For every observed paired interaction 30 observations are generated from the model, i.e.: E[Y = bootstrap2 | X = bootstrap1 ]. The second bootstrap dataset is sparse in interactions with respect to first in order to mimic different coverage. The X axis is log transformed.

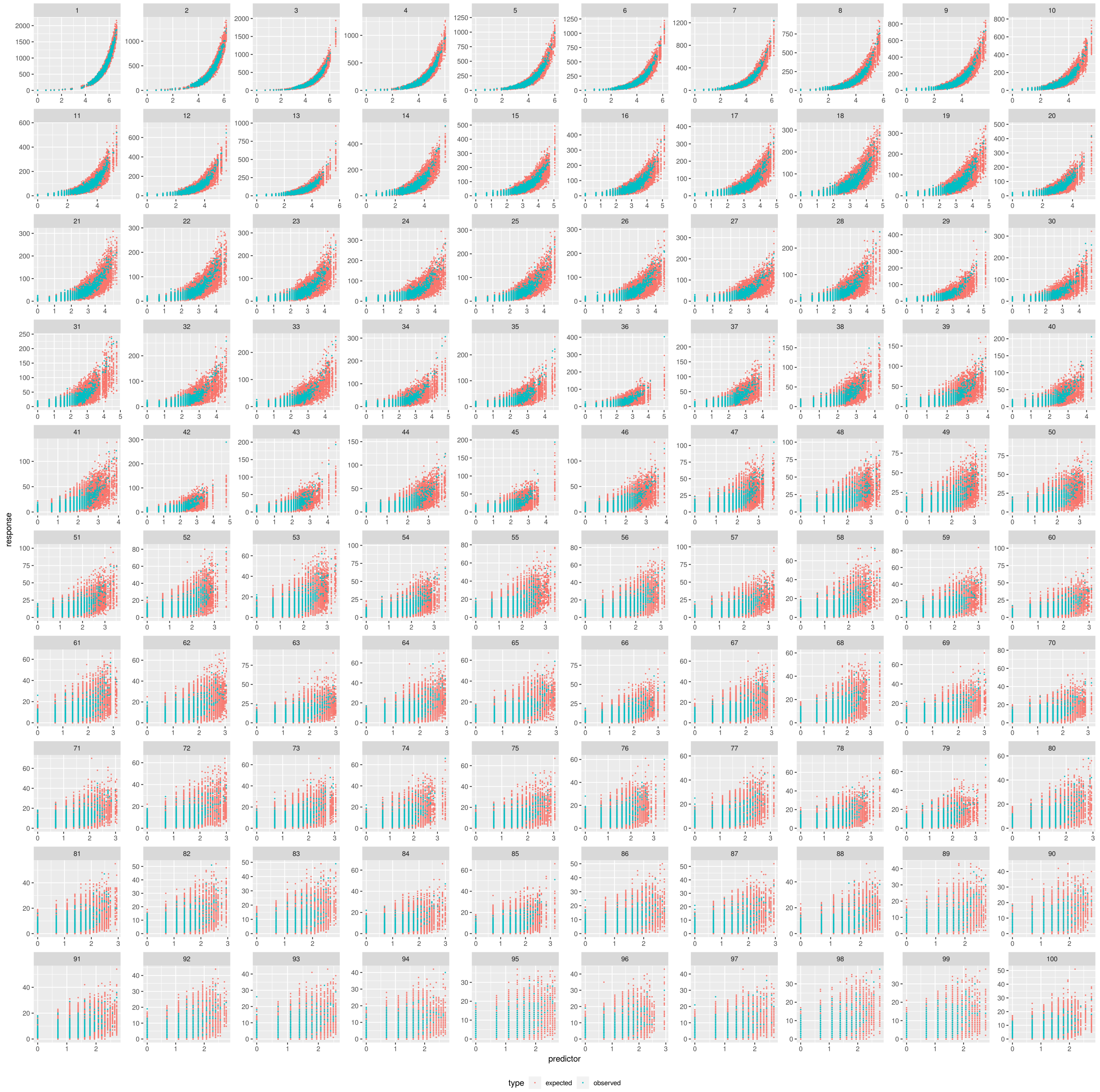


Figure 12: **Expected vs observed paired interactions for IMR90 bootstrapped data.** The same as in fig. 11, but now: E[Y = bootstrap1 | X = bootstrap2].

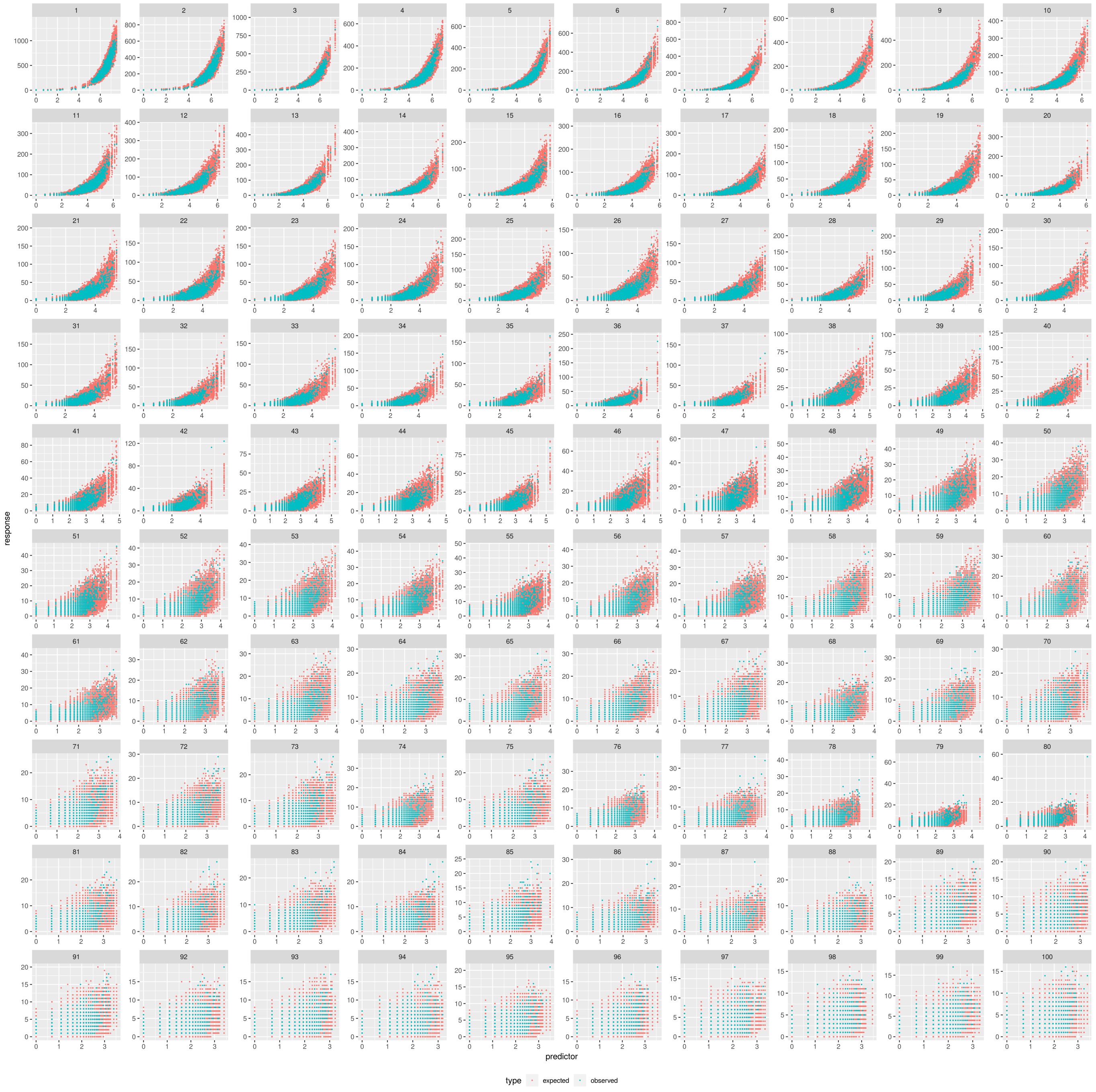


Figure 13: **Expected vs observed paired interactions for IMR90 biological replicates data.** The same as in fig. 11, but now: E[Y = replicate2 | X = replicate1].

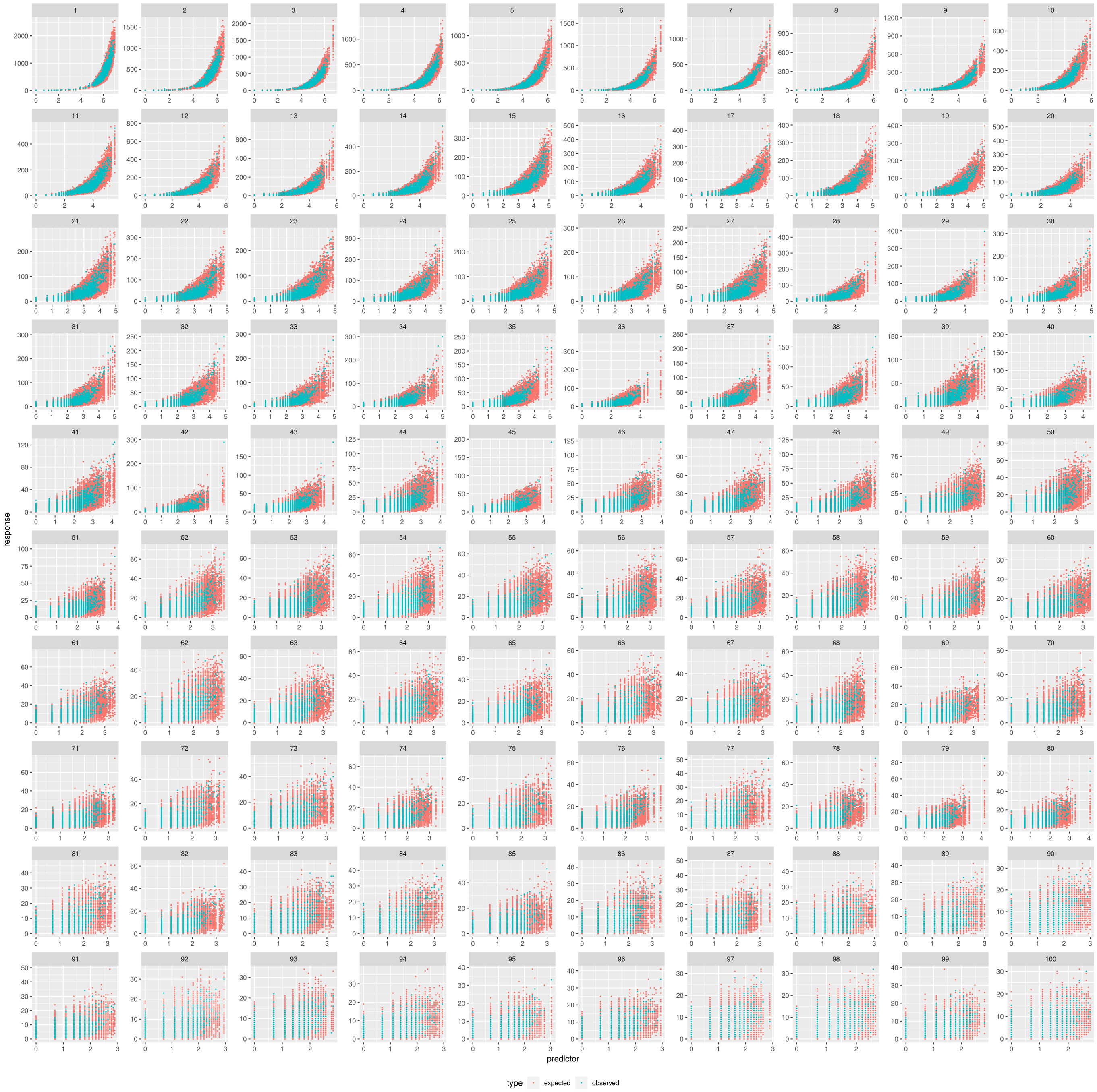


Figure 14: **Expected vs observed paired interactions for IMR90 biological replicates data.** The same as in fig. 11, but now: E[Y = replicate1 | X = replicate2].

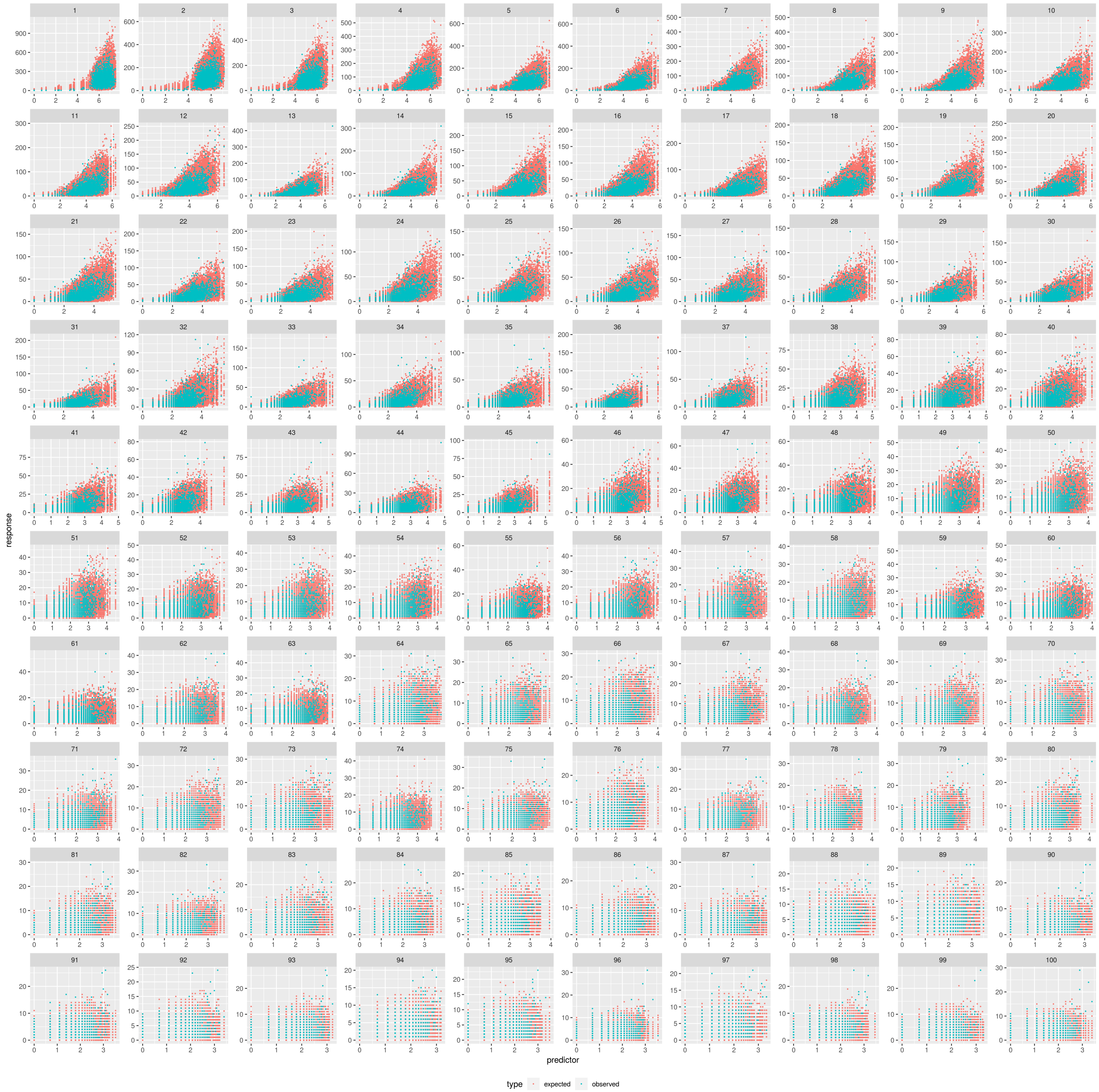


Figure 15: **Expected vs observed paired interactions for IMR90 vs MSC data.** The same as in fig. 11, but now: E[Y = MSC | X = IMR90].

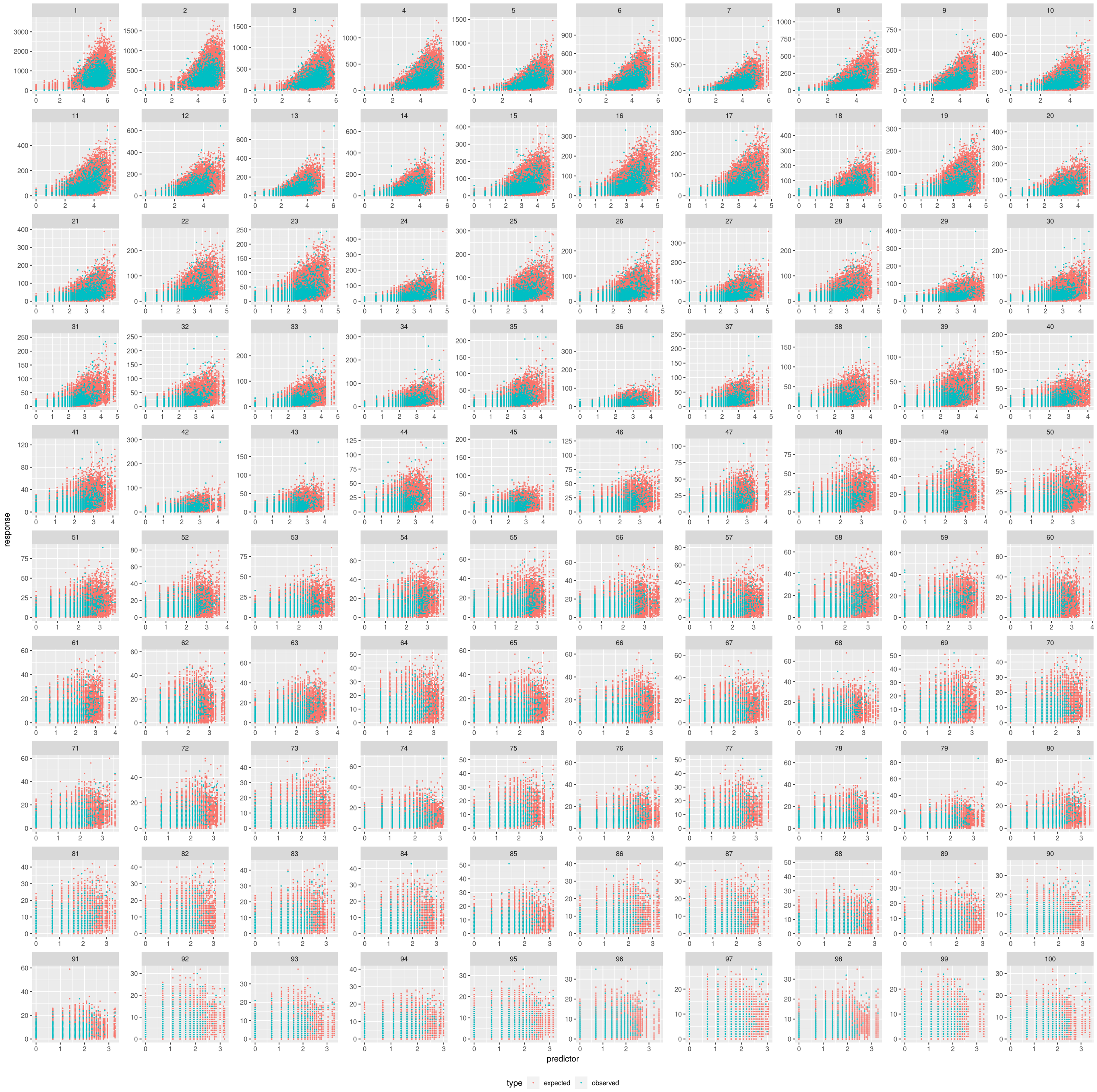


Figure 16: **Expected vs observed paired interactions for IMR90 vs MSC data.** The same as in fig. 11, but now: E[Y = IMR90 | X = MSC].

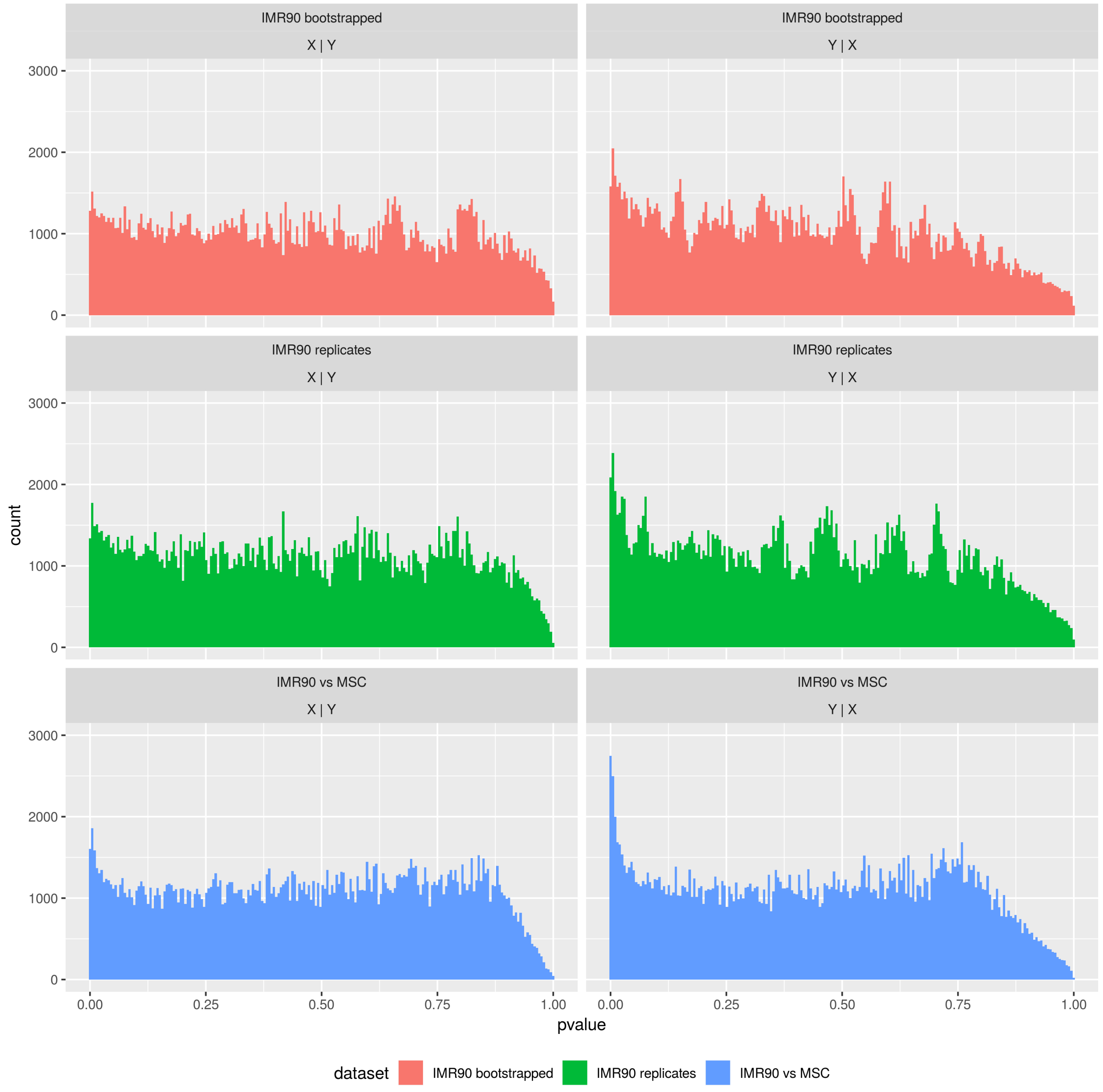


Figure 17: **P-value distribution for IMR90 bootstrapped, IMR90 biological replicates and IMR90 vs MSC.**

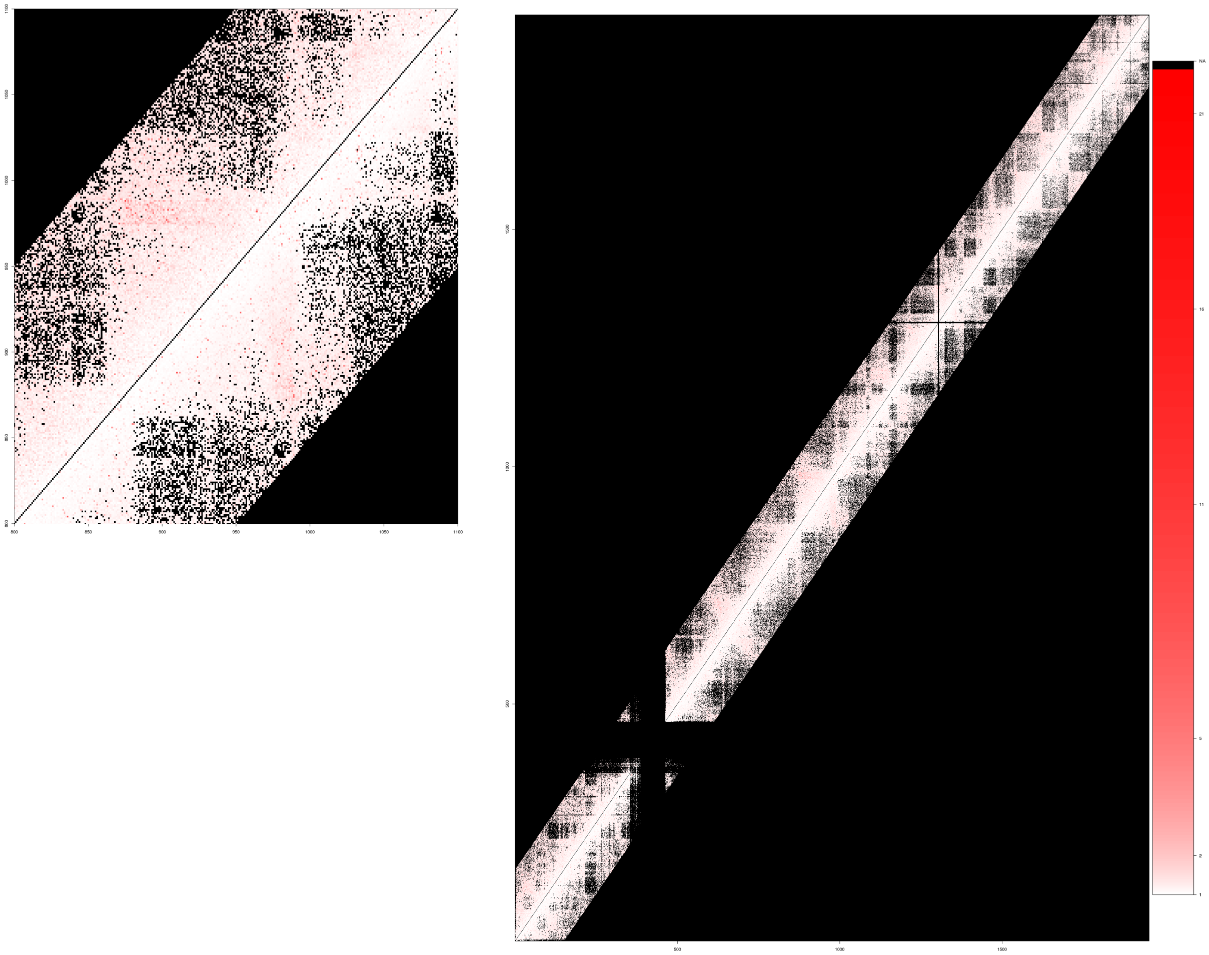

Figure 18: **Significance/Differential map of IMR90 bootstrap 1 vs 2.** Each cell intensity equals $-log(\text{p})$ of enrichment of 2 with respect to 1.

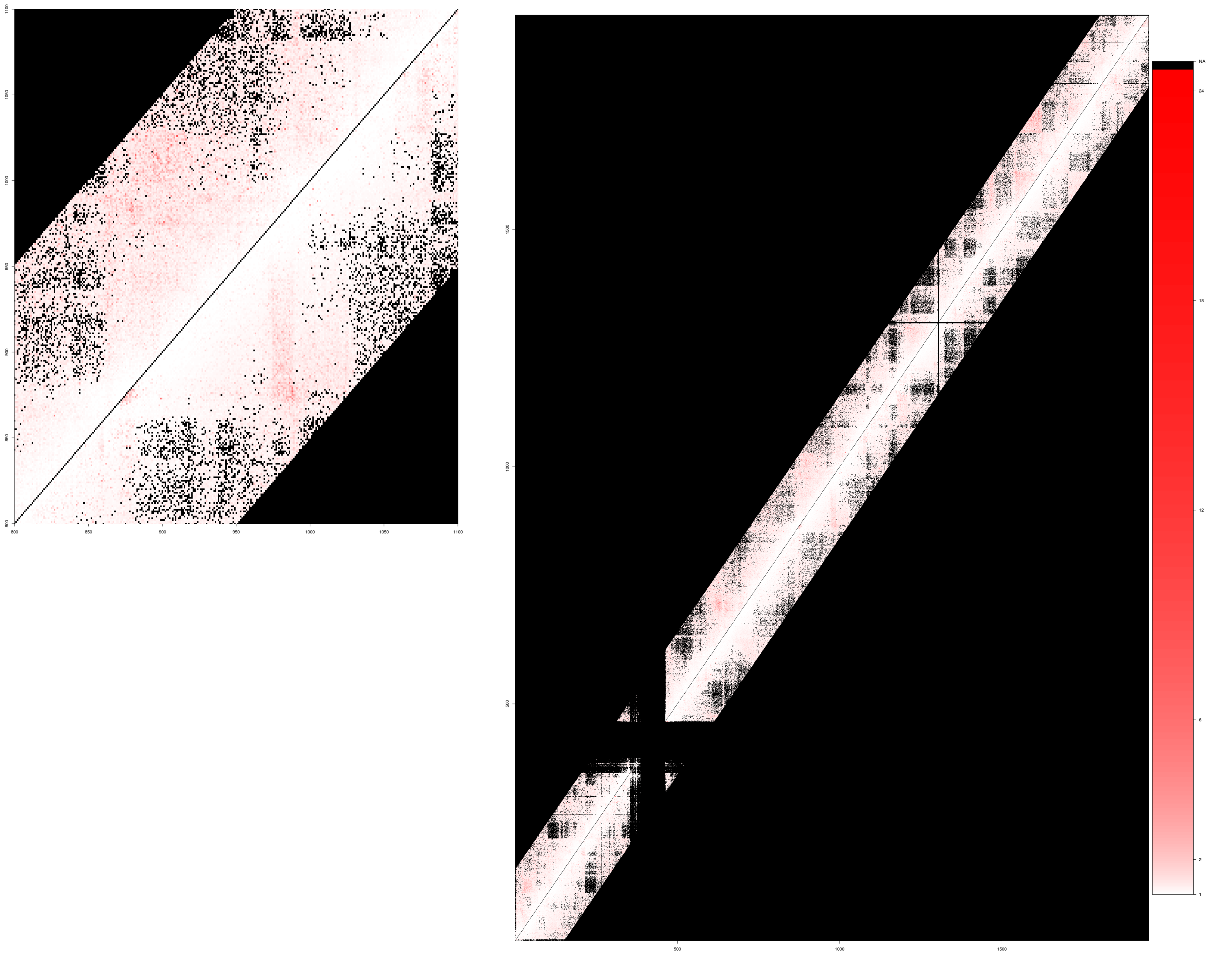

Figure 19: **Significance/Differential map of IMR90 biological replicate 1 vs 2.** The same as in fig. 18, but different Hi-C datasets are compared.

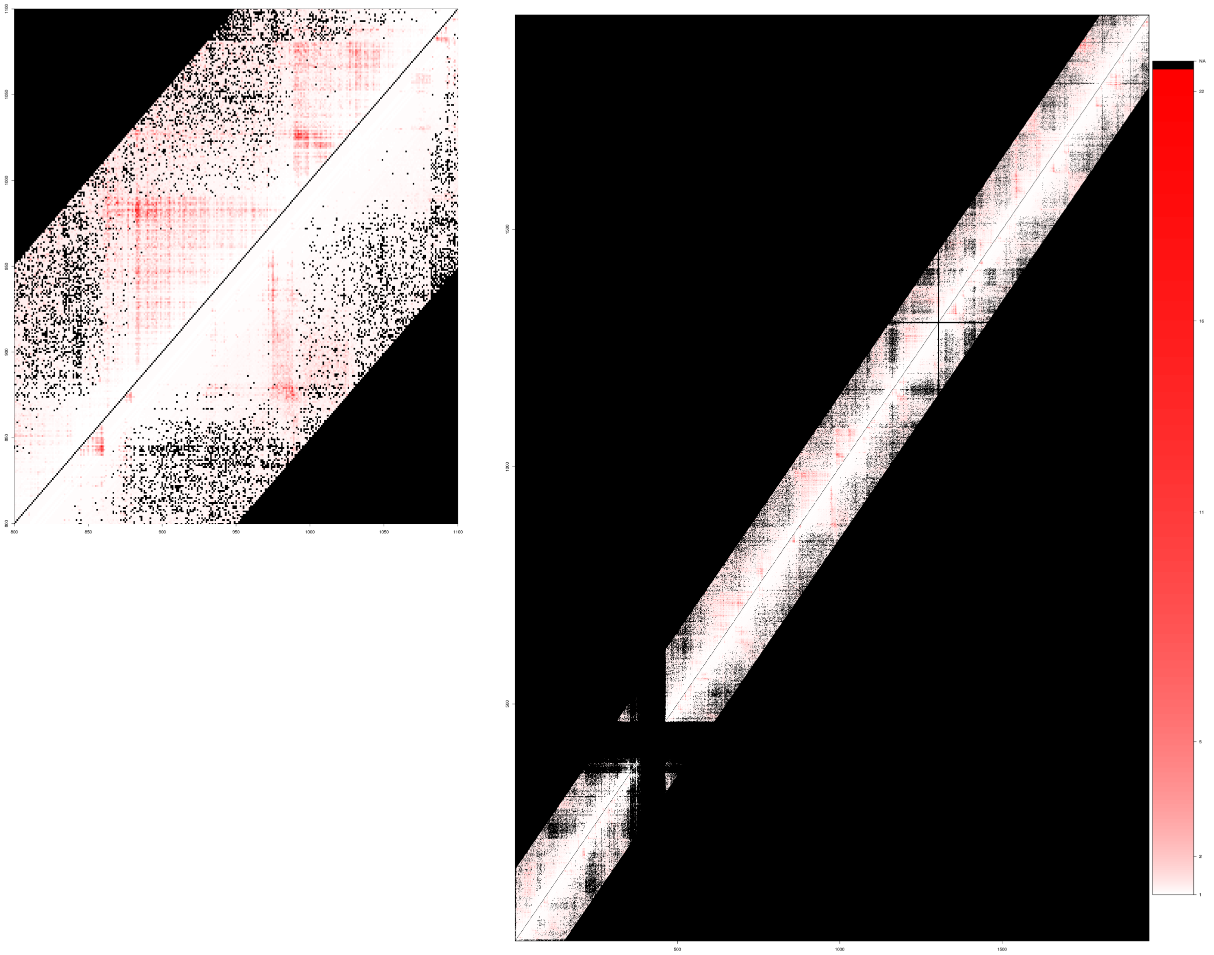

Figure 20: **Significance/Differential map of IMR90 vs MSC.** The same as in fig. 18, but different Hi-C datasets are compared.

a)

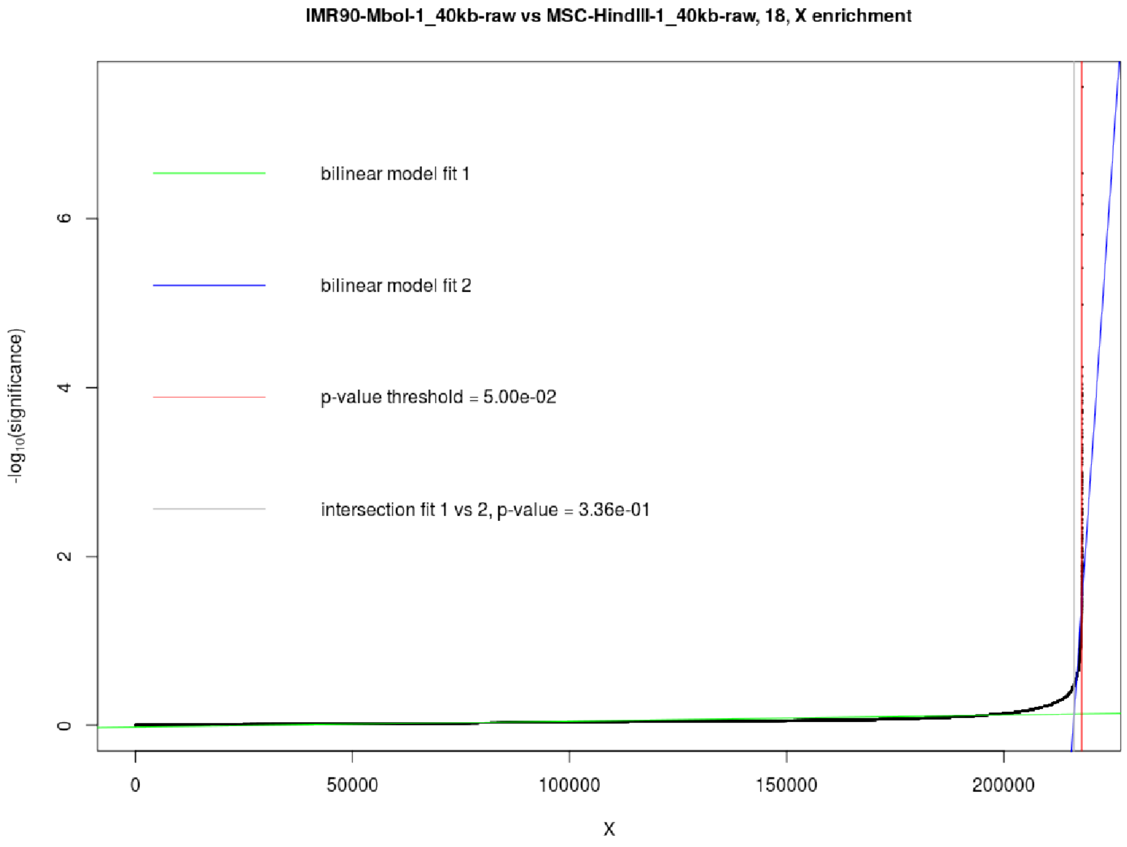


b)

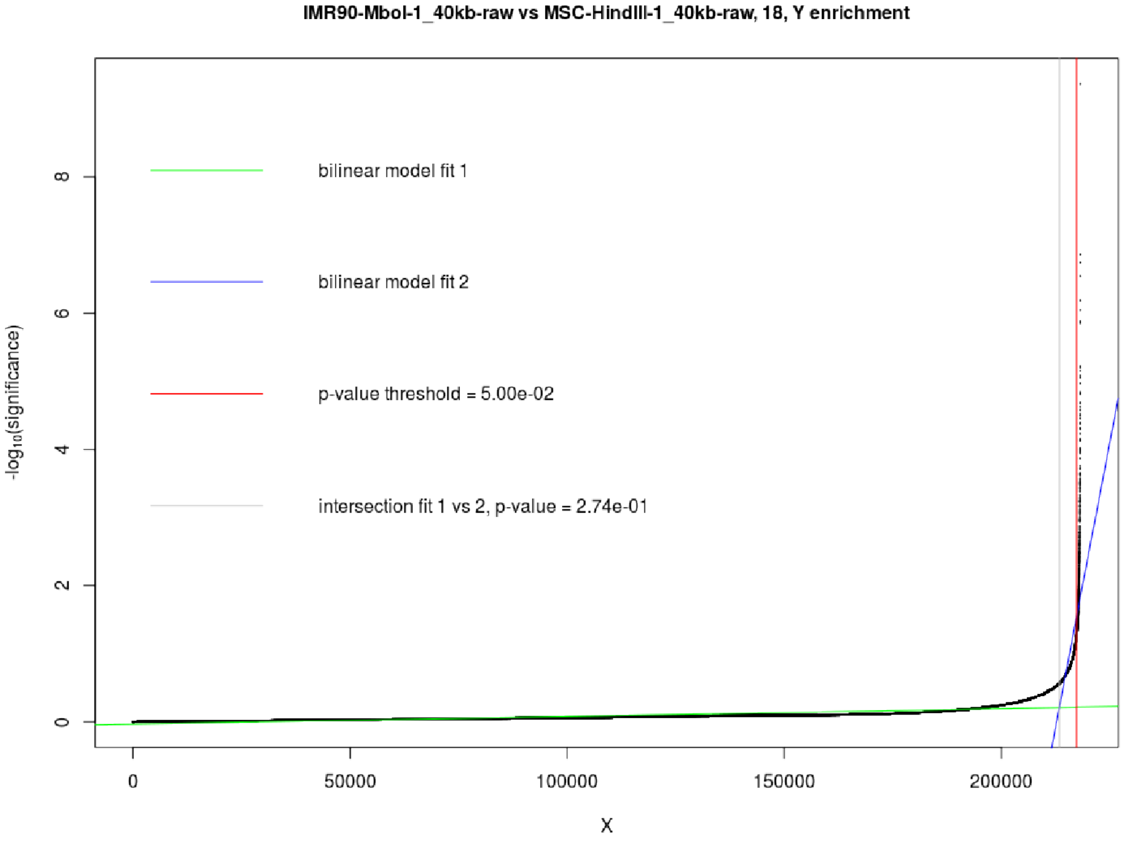


Figure 21: **Detection of Long Range Differentially Interacting Regions, bilinear model fit.**

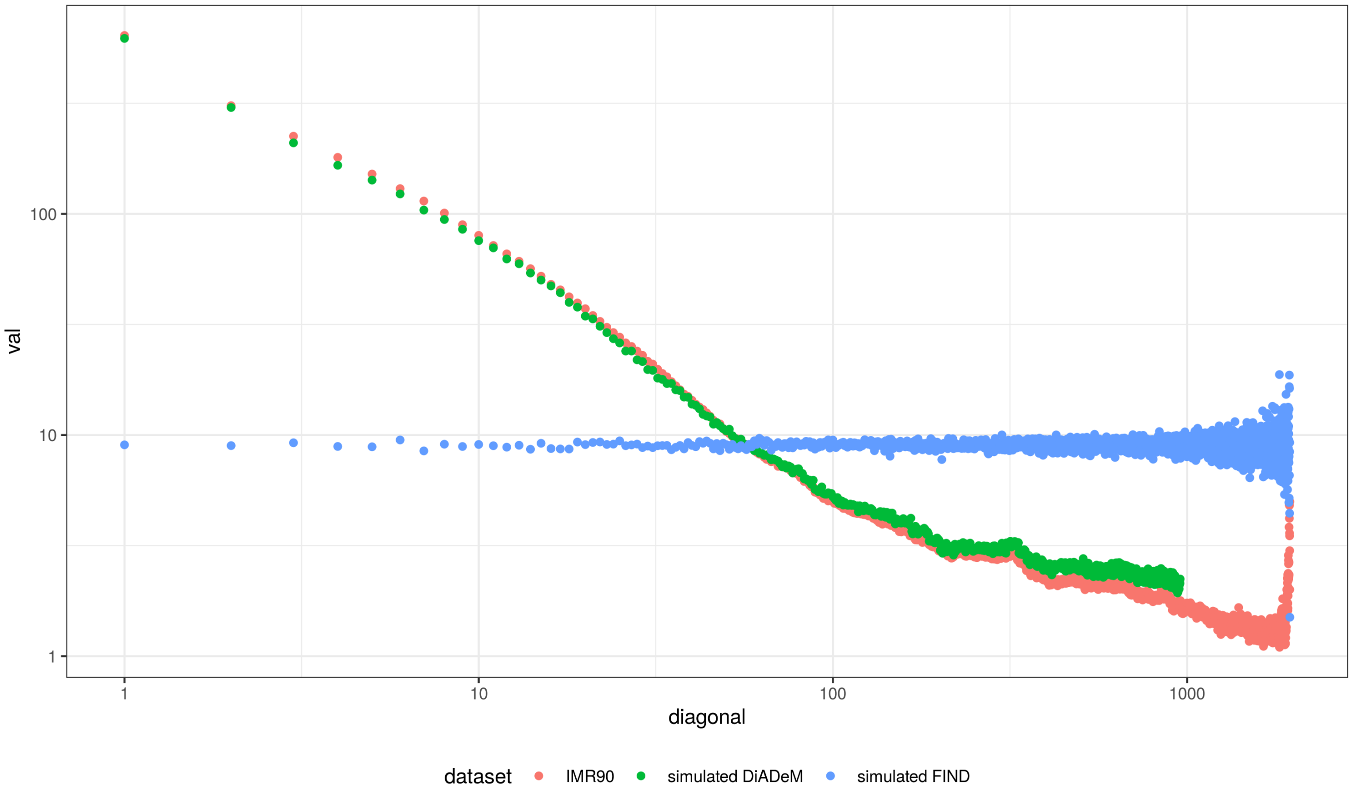


Figure 22: **Comparison of decays for real Hi-C IMR90 dataset, and artificial dataset simulated from IMR90 - by 2 approaches: DiADeM and FIND.**

### 2 Supplementary tables

Table 1: Diagonal-wise interaction correlations

| study1 | cell1 | study2 | cell2 | chromosome | coverage1 | coverage2 | number.significant | fraction.significant |
|---|---|---|---|---|---|---|---|---|
| GSE87112-Schmitt-2016 | PO3-HindIII-1 | GSE87112-Schmitt-2016 | PA3-HindIII-1 | 1 | 102.52 | 127.34 | 243 | 0.39 |
| GSE87112-Schmitt-2016 | LV-HindIII-1 | GSE87112-Schmitt-2016 | PO3-HindIII-1 | 1 | 1,835.39 | 102.52 | 88 | 0.14 |
| GSE87112-Schmitt-2016 | LV-HindIII-1 | GSE87112-Schmitt-2016 | PA3-HindIII-1 | 1 | 1,835.39 | 127.34 | 106 | 0.17 |
| GSE87112-Schmitt-2016 | LV-HindIII-1 | GSE52457-Dixon-2015 | MDC-HindIII-2 | 1 | 1,835.39 | 1,993.82 | 244 | 0.39 |
| GSE87112-Schmitt-2016 | LV-HindIII-1 | GSE52457-Dixon-2015 | MDC-HindIII-1 | 1 | 1,835.39 | 2,114.95 | 238 | 0.38 |
| GSE52457-Dixon-2015 | MDC-HindIII-2 | GSE87112-Schmitt-2016 | PO3-HindIII-1 | 1 | 1,993.82 | 102.52 | 81 | 0.13 |
| GSE52457-Dixon-2015 | MDC-HindIII-2 | GSE87112-Schmitt-2016 | PA3-HindIII-1 | 1 | 1,993.82 | 127.34 | 106 | 0.17 |
| GSE52457-Dixon-2015 | MDC-HindIII-1 | GSE87112-Schmitt-2016 | PO3-HindIII-1 | 1 | 2,114.95 | 102.52 | 85 | 0.14 |
| GSE52457-Dixon-2015 | MDC-HindIII-1 | GSE87112-Schmitt-2016 | PA3-HindIII-1 | 1 | 2,114.95 | 127.34 | 109 | 0.17 |
| GSE87112-Schmitt-2016 | H1-HindIII-2 | GSE87112-Schmitt-2016 | PO3-HindIII-1 | 1 | 2,230.06 | 102.52 | 96 | 0.15 |
| GSE87112-Schmitt-2016 | H1-HindIII-2 | GSE87112-Schmitt-2016 | PA3-HindIII-1 | 1 | 2,230.06 | 127.34 | 116 | 0.19 |
| GSE87112-Schmitt-2016 | H1-HindIII-2 | GSE87112-Schmitt-2016 | LV-HindIII-1 | 1 | 2,230.06 | 1,835.39 | 339 | 0.54 |
| GSE87112-Schmitt-2016 | H1-HindIII-2 | GSE52457-Dixon-2015 | MDC-HindIII-2 | 1 | 2,230.06 | 1,993.82 | 549 | 0.88 |
| GSE87112-Schmitt-2016 | H1-HindIII-2 | GSE52457-Dixon-2015 | MDC-HindIII-1 | 1 | 2,230.06 | 2,114.95 | 610 | 0.98 |
| GSE63525-Rao-2014 | IMR90-MboI-2 | GSE87112-Schmitt-2016 | PO3-HindIII-1 | 1 | 3,182.77 | 102.52 | 70 | 0.11 |
| GSE63525-Rao-2014 | IMR90-MboI-2 | GSE87112-Schmitt-2016 | PA3-HindIII-1 | 1 | 3,182.77 | 127.34 | 93 | 0.15 |
| GSE63525-Rao-2014 | IMR90-MboI-2 | GSE87112-Schmitt-2016 | LV-HindIII-1 | 1 | 3,182.77 | 1,835.39 | 160 | 0.26 |
| GSE63525-Rao-2014 | IMR90-MboI-2 | GSE52457-Dixon-2015 | MDC-HindIII-2 | 1 | 3,182.77 | 1,993.82 | 228 | 0.37 |
| GSE63525-Rao-2014 | IMR90-MboI-2 | GSE52457-Dixon-2015 | MDC-HindIII-1 | 1 | 3,182.77 | 2,114.95 | 242 | 0.39 |
| GSE63525-Rao-2014 | IMR90-MboI-2 | GSE87112-Schmitt-2016 | H1-HindIII-2 | 1 | 3,182.77 | 2,230.06 | 220 | 0.35 |
| GSE63525-Rao-2014 | IMR90-MboI-2 | GSE87112-Schmitt-2016 | H1-HindIII-1 | 1 | 3,182.77 | 3,601.07 | 267 | 0.43 |
| GSE87112-Schmitt-2016 | H1-HindIII-1 | GSE87112-Schmitt-2016 | PO3-HindIII-1 | 1 | 3,601.07 | 102.52 | 103 | 0.17 |
| GSE87112-Schmitt-2016 | H1-HindIII-1 | GSE87112-Schmitt-2016 | PA3-HindIII-1 | 1 | 3,601.07 | 127.34 | 132 | 0.21 |
| GSE87112-Schmitt-2016 | H1-HindIII-1 | GSE87112-Schmitt-2016 | LV-HindIII-1 | 1 | 3,601.07 | 1,835.39 | 453 | 0.73 |
| GSE87112-Schmitt-2016 | H1-HindIII-1 | GSE52457-Dixon-2015 | MDC-HindIII-2 | 1 | 3,601.07 | 1,993.82 | 613 | 0.98 |
| GSE87112-Schmitt-2016 | H1-HindIII-1 | GSE52457-Dixon-2015 | MDC-HindIII-1 | 1 | 3,601.07 | 2,114.95 | 620 | 1 |
| GSE63525-Rao-2014 | IMR90-MboI-1 | GSE87112-Schmitt-2016 | PO3-HindIII-1 | 1 | 5,408.54 | 102.52 | 83 | 0.13 |
| GSE63525-Rao-2014 | IMR90-MboI-1 | GSE87112-Schmitt-2016 | PA3-HindIII-1 | 1 | 5,408.54 | 127.34 | 109 | 0.17 |
| GSE63525-Rao-2014 | IMR90-MboI-1 | GSE87112-Schmitt-2016 | LV-HindIII-1 | 1 | 5,408.54 | 1,835.39 | 175 | 0.28 |
| GSE63525-Rao-2014 | IMR90-MboI-1 | GSE52457-Dixon-2015 | MDC-HindIII-2 | 1 | 5,408.54 | 1,993.82 | 220 | 0.35 |
| GSE63525-Rao-2014 | IMR90-MboI-1 | GSE52457-Dixon-2015 | MDC-HindIII-1 | 1 | 5,408.54 | 2,114.95 | 258 | 0.41 |
| GSE63525-Rao-2014 | IMR90-MboI-1 | GSE87112-Schmitt-2016 | H1-HindIII-2 | 1 | 5,408.54 | 2,230.06 | 197 | 0.32 |
| GSE63525-Rao-2014 | IMR90-MboI-1 | GSE87112-Schmitt-2016 | H1-HindIII-1 | 1 | 5,408.54 | 3,601.07 | 242 | 0.39 |
| GSE87112-Schmitt-2016 | PO3-HindIII-1 | GSE87112-Schmitt-2016 | PA3-HindIII-1 | 10 | 107.59 | 132.66 | 128 | 0.38 |
| GSE87112-Schmitt-2016 | LV-HindIII-1 | GSE87112-Schmitt-2016 | PO3-HindIII-1 | 10 | 1,900.21 | 107.59 | 66 | 0.19 |
| GSE87112-Schmitt-2016 | LV-HindIII-1 | GSE87112-Schmitt-2016 | PA3-HindIII-1 | 10 | 1,900.21 | 132.66 | 90 | 0.27 |
| GSE87112-Schmitt-2016 | LV-HindIII-1 | GSE52457-Dixon-2015 | MDC-HindIII-2 | 10 | 1,900.21 | 2,099.26 | 177 | 0.52 |
| GSE87112-Schmitt-2016 | LV-HindIII-1 | GSE52457-Dixon-2015 | MDC-HindIII-1 | 10 | 1,900.21 | 2,188.23 | 178 | 0.53 |
| GSE52457-Dixon-2015 | MDC-HindIII-2 | GSE87112-Schmitt-2016 | PO3-HindIII-1 | 10 | 2,099.26 | 107.59 | 68 | 0.2 |
| GSE52457-Dixon-2015 | MDC-HindIII-2 | GSE87112-Schmitt-2016 | PA3-HindIII-1 | 10 | 2,099.26 | 132.66 | 87 | 0.26 |
| GSE87112-Schmitt-2016 | H1-HindIII-2 | GSE87112-Schmitt-2016 | PO3-HindIII-1 | 10 | 2,187.22 | 107.59 | 75 | 0.22 |
| GSE87112-Schmitt-2016 | H1-HindIII-2 | GSE87112-Schmitt-2016 | PA3-HindIII-1 | 10 | 2,187.22 | 132.66 | 102 | 0.3 |
| GSE87112-Schmitt-2016 | H1-HindIII-2 | GSE87112-Schmitt-2016 | LV-HindIII-1 | 10 | 2,187.22 | 1,900.21 | 216 | 0.64 |
| GSE87112-Schmitt-2016 | H1-HindIII-2 | GSE52457-Dixon-2015 | MDC-HindIII-2 | 10 | 2,187.22 | 2,099.26 | 302 | 0.89 |
| GSE87112-Schmitt-2016 | H1-HindIII-2 | GSE52457-Dixon-2015 | MDC-HindIII-1 | 10 | 2,187.22 | 2,188.23 | 321 | 0.95 |
| GSE52457-Dixon-2015 | MDC-HindIII-1 | GSE87112-Schmitt-2016 | PO3-HindIII-1 | 10 | 2,188.23 | 107.59 | 72 | 0.21 |
| GSE52457-Dixon-2015 | MDC-HindIII-1 | GSE87112-Schmitt-2016 | PA3-HindIII-1 | 10 | 2,188.23 | 132.66 | 89 | 0.26 |
| GSE63525-Rao-2014 | IMR90-MboI-2 | GSE87112-Schmitt-2016 | PO3-HindIII-1 | 10 | 3,258.35 | 107.59 | 67 | 0.2 |
| GSE63525-Rao-2014 | IMR90-MboI-2 | GSE87112-Schmitt-2016 | PA3-HindIII-1 | 10 | 3,258.35 | 132.66 | 80 | 0.24 |
| GSE63525-Rao-2014 | IMR90-MboI-2 | GSE87112-Schmitt-2016 | LV-HindIII-1 | 10 | 3,258.35 | 1,900.21 | 159 | 0.47 |
| GSE63525-Rao-2014 | IMR90-MboI-2 | GSE52457-Dixon-2015 | MDC-HindIII-2 | 10 | 3,258.35 | 2,099.26 | 181 | 0.53 |
| GSE63525-Rao-2014 | IMR90-MboI-2 | GSE87112-Schmitt-2016 | H1-HindIII-2 | 10 | 3,258.35 | 2,187.22 | 189 | 0.56 |
| GSE63525-Rao-2014 | IMR90-MboI-2 | GSE52457-Dixon-2015 | MDC-HindIII-1 | 10 | 3,258.35 | 2,188.23 | 188 | 0.55 |
| GSE63525-Rao-2014 | IMR90-MboI-2 | GSE87112-Schmitt-2016 | H1-HindIII-1 | 10 | 3,258.35 | 3,593.06 | 214 | 0.63 |
| GSE87112-Schmitt-2016 | H1-HindIII-1 | GSE87112-Schmitt-2016 | PO3-HindIII-1 | 10 | 3,593.06 | 107.59 | 84 | 0.25 |
| GSE87112-Schmitt-2016 | H1-HindIII-1 | GSE87112-Schmitt-2016 | PA3-HindIII-1 | 10 | 3,593.06 | 132.66 | 111 | 0.33 |
| GSE87112-Schmitt-2016 | H1-HindIII-1 | GSE87112-Schmitt-2016 | LV-HindIII-1 | 10 | 3,593.06 | 1,900.21 | 257 | 0.76 |
| GSE87112-Schmitt-2016 | H1-HindIII-1 | GSE52457-Dixon-2015 | MDC-HindIII-2 | 10 | 3,593.06 | 2,099.26 | 327 | 0.96 |
| GSE87112-Schmitt-2016 | H1-HindIII-1 | GSE52457-Dixon-2015 | MDC-HindIII-1 | 10 | 3,593.06 | 2,188.23 | 336 | 0.99 |
| GSE63525-Rao-2014 | IMR90-MboI-1 | GSE87112-Schmitt-2016 | PO3-HindIII-1 | 10 | 5,522.43 | 107.59 | 69 | 0.2 |
| GSE63525-Rao-2014 | IMR90-MboI-1 | GSE87112-Schmitt-2016 | PA3-HindIII-1 | 10 | 5,522.43 | 132.66 | 93 | 0.27 |
| GSE63525-Rao-2014 | IMR90-MboI-1 | GSE87112-Schmitt-2016 | LV-HindIII-1 | 10 | 5,522.43 | 1,900.21 | 164 | 0.48 |
| GSE63525-Rao-2014 | IMR90-MboI-1 | GSE52457-Dixon-2015 | MDC-HindIII-2 | 10 | 5,522.43 | 2,099.26 | 186 | 0.55 |
| GSE63525-Rao-2014 | IMR90-MboI-1 | GSE87112-Schmitt-2016 | H1-HindIII-2 | 10 | 5,522.43 | 2,187.22 | 183 | 0.54 |
| GSE63525-Rao-2014 | IMR90-MboI-1 | GSE52457-Dixon-2015 | MDC-HindIII-1 | 10 | 5,522.43 | 2,188.23 | 197 | 0.58 |
| GSE63525-Rao-2014 | IMR90-MboI-1 | GSE87112-Schmitt-2016 | H1-HindIII-1 | 10 | 5,522.43 | 3,593.06 | 211 | 0.62 |
| GSE87112-Schmitt-2016 | PO3-HindIII-1 | GSE87112-Schmitt-2016 | PA3-HindIII-1 | 18 | 97.69 | 127.7 | 49 | 0.25 |
| GSE87112-Schmitt-2016 | LV-HindIII-1 | GSE87112-Schmitt-2016 | PO3-HindIII-1 | 18 | 1,970.51 | 97.69 | 53 | 0.27 |
| GSE87112-Schmitt-2016 | LV-HindIII-1 | GSE87112-Schmitt-2016 | PA3-HindIII-1 | 18 | 1,970.51 | 127.7 | 60 | 0.31 |
| GSE87112-Schmitt-2016 | LV-HindIII-1 | GSE52457-Dixon-2015 | MDC-HindIII-2 | 18 | 1,970.51 | 2,382.45 | 152 | 0.78 |
| GSE87112-Schmitt-2016 | LV-HindIII-1 | GSE52457-Dixon-2015 | MDC-HindIII-1 | 18 | 1,970.51 | 2,393.13 | 141 | 0.72 |
| GSE87112-Schmitt-2016 | H1-HindIII-2 | GSE87112-Schmitt-2016 | PO3-HindIII-1 | 18 | 2,297.91 | 97.69 | 52 | 0.27 |
| GSE87112-Schmitt-2016 | H1-HindIII-2 | GSE87112-Schmitt-2016 | PA3-HindIII-1 | 18 | 2,297.91 | 127.7 | 70 | 0.36 |
| GSE87112-Schmitt-2016 | H1-HindIII-2 | GSE87112-Schmitt-2016 | LV-HindIII-1 | 18 | 2,297.91 | 1,970.51 | 156 | 0.8 |
| GSE87112-Schmitt-2016 | H1-HindIII-2 | GSE52457-Dixon-2015 | MDC-HindIII-2 | 18 | 2,297.91 | 2,382.45 | 186 | 0.95 |
| GSE87112-Schmitt-2016 | H1-HindIII-2 | GSE52457-Dixon-2015 | MDC-HindIII-1 | 18 | 2,297.91 | 2,393.13 | 193 | 0.99 |
| GSE52457-Dixon-2015 | MDC-HindIII-2 | GSE87112-Schmitt-2016 | PO3-HindIII-1 | 18 | 2,382.45 | 97.69 | 53 | 0.27 |
| GSE52457-Dixon-2015 | MDC-HindIII-2 | GSE87112-Schmitt-2016 | PA3-HindIII-1 | 18 | 2,382.45 | 127.7 | 75 | 0.38 |
| GSE52457-Dixon-2015 | MDC-HindIII-1 | GSE87112-Schmitt-2016 | PO3-HindIII-1 | 18 | 2,393.13 | 97.69 | 54 | 0.28 |
| GSE52457-Dixon-2015 | MDC-HindIII-1 | GSE87112-Schmitt-2016 | PA3-HindIII-1 | 18 | 2,393.13 | 127.7 | 69 | 0.35 |
| GSE63525-Rao-2014 | IMR90-MboI-2 | GSE87112-Schmitt-2016 | PO3-HindIII-1 | 18 | 3,268.8 | 97.69 | 41 | 0.21 |
| GSE63525-Rao-2014 | IMR90-MboI-2 | GSE87112-Schmitt-2016 | PA3-HindIII-1 | 18 | 3,268.8 | 127.7 | 54 | 0.28 |
| GSE63525-Rao-2014 | IMR90-MboI-2 | GSE87112-Schmitt-2016 | LV-HindIII-1 | 18 | 3,268.8 | 1,970.51 | 134 | 0.69 |
| GSE63525-Rao-2014 | IMR90-MboI-2 | GSE87112-Schmitt-2016 | H1-HindIII-2 | 18 | 3,268.8 | 2,297.91 | 132 | 0.68 |
| GSE63525-Rao-2014 | IMR90-MboI-2 | GSE52457-Dixon-2015 | MDC-HindIII-2 | 18 | 3,268.8 | 2,382.45 | 122 | 0.63 |
| GSE63525-Rao-2014 | IMR90-MboI-2 | GSE52457-Dixon-2015 | MDC-HindIII-1 | 18 | 3,268.8 | 2,393.13 | 125 | 0.64 |
| GSE63525-Rao-2014 | IMR90-MboI-2 | GSE87112-Schmitt-2016 | H1-HindIII-1 | 18 | 3,268.8 | 3,864.66 | 152 | 0.78 |
| GSE87112-Schmitt-2016 | H1-HindIII-1 | GSE87112-Schmitt-2016 | PO3-HindIII-1 | 18 | 3,864.66 | 97.69 | 58 | 0.3 |
| GSE87112-Schmitt-2016 | H1-HindIII-1 | GSE87112-Schmitt-2016 | PA3-HindIII-1 | 18 | 3,864.66 | 127.7 | 68 | 0.35 |
| GSE87112-Schmitt-2016 | H1-HindIII-1 | GSE87112-Schmitt-2016 | LV-HindIII-1 | 18 | 3,864.66 | 1,970.51 | 181 | 0.93 |
| GSE87112-Schmitt-2016 | H1-HindIII-1 | GSE52457-Dixon-2015 | MDC-HindIII-2 | 18 | 3,864.66 | 2,382.45 | 194 | 0.99 |
| GSE87112-Schmitt-2016 | H1-HindIII-1 | GSE52457-Dixon-2015 | MDC-HindIII-1 | 18 | 3,864.66 | 2,393.13 | 195 | 1 |
| GSE63525-Rao-2014 | IMR90-MboI-1 | GSE87112-Schmitt-2016 | PO3-HindIII-1 | 18 | 5,261.43 | 97.69 | 48 | 0.25 |
| GSE63525-Rao-2014 | IMR90-MboI-1 | GSE87112-Schmitt-2016 | PA3-HindIII-1 | 18 | 5,261.43 | 127.7 | 62 | 0.32 |
| GSE63525-Rao-2014 | IMR90-MboI-1 | GSE87112-Schmitt-2016 | LV-HindIII-1 | 18 | 5,261.43 | 1,970.51 | 136 | 0.7 |
| GSE63525-Rao-2014 | IMR90-MboI-1 | GSE87112-Schmitt-2016 | H1-HindIII-2 | 18 | 5,261.43 | 2,297.91 | 117 | 0.6 |
| GSE63525-Rao-2014 | IMR90-MboI-1 | GSE52457-Dixon-2015 | MDC-HindIII-2 | 18 | 5,261.43 | 2,382.45 | 120 | 0.62 |
| GSE63525-Rao-2014 | IMR90-MboI-1 | GSE52457-Dixon-2015 | MDC-HindIII-1 | 18 | 5,261.43 | 2,393.13 | 127 | 0.65 |
| GSE63525-Rao-2014 | IMR90-MboI-1 | GSE87112-Schmitt-2016 | H1-HindIII-1 | 18 | 5,261.43 | 3,864.66 | 135 | 0.69 |
| GSE52457-Dixon-2015 | MDC-HindIII-1 | GSE52457-Dixon-2015 | MDC-HindIII-2 | 1 | 2,114.95 | 1,993.82 | 502 | 0.81 |
| GSE87112-Schmitt-2016 | H1-HindIII-1 | GSE87112-Schmitt-2016 | H1-HindIII-2 | 1 | 3,601.07 | 2,230.06 | 623 | 1 |
| GSE63525-Rao-2014 | IMR90-MboI-1 | GSE63525-Rao-2014 | IMR90-MboI-2 | 1 | 5,408.54 | 3,182.77 | 623 | 1 |
| GSE52457-Dixon-2015 | MDC-HindIII-1 | GSE52457-Dixon-2015 | MDC-HindIII-2 | 10 | 2,188.23 | 2,099.26 | 286 | 0.84 |
| GSE87112-Schmitt-2016 | H1-HindIII-1 | GSE87112-Schmitt-2016 | H1-HindIII-2 | 10 | 3,593.06 | 2,187.22 | 339 | 1 |

| | | | | | | | | |
|---|---|---|---|---|---|---|---|---|
| GSE63525-Rao-2014 | IMR90-MboI-1 | GSE63525-Rao-2014 | IMR90-MboI-2 | 10 | 5,522.43 | 3,258.35 | 339 | 1 |
| GSE52457-Dixon-2015 | MDC-HindIII-1 | GSE52457-Dixon-2015 | MDC-HindIII-2 | 18 | 2,393.13 | 2,382.45 | 189 | 0.97 |
| GSE87112-Schmitt-2016 | H1-HindIII-1 | GSE87112-Schmitt-2016 | H1-HindIII-2 | 18 | 3,864.66 | 2,297.91 | 195 | 1 |
| GSE63525-Rao-2014 | IMR90-MboI-1 | GSE63525-Rao-2014 | IMR90-MboI-2 | 18 | 5,261.43 | 3,268.8 | 195 | 1 |

Table 2: Cumulative fraction of contacts in cumulative fraction of diagonals

| study | cell | chromosome | first 5 | first 10 | first 15 | first 20 | first 30 | first 40 | first 50 |
|---|---|---|---|---|---|---|---|---|---|
| GSE63525-Rao-2014 | IMR90-MboI-1 | 1 | 0.83 | 0.89 | 0.92 | 0.94 | 0.96 | 0.97 | 0.98 |
| GSE63525-Rao-2014 | IMR90-MboI-1 | 10 | 0.81 | 0.86 | 0.89 | 0.92 | 0.95 | 0.97 | 0.98 |
| GSE63525-Rao-2014 | IMR90-MboI-1 | 18 | 0.78 | 0.83 | 0.86 | 0.89 | 0.93 | 0.96 | 0.98 |
| GSE63525-Rao-2014 | IMR90-MboI-2 | 1 | 0.83 | 0.89 | 0.92 | 0.94 | 0.96 | 0.97 | 0.98 |
| GSE63525-Rao-2014 | IMR90-MboI-2 | 10 | 0.81 | 0.86 | 0.89 | 0.92 | 0.95 | 0.97 | 0.98 |
| GSE63525-Rao-2014 | IMR90-MboI-2 | 18 | 0.78 | 0.83 | 0.86 | 0.89 | 0.93 | 0.96 | 0.98 |
| GSE87112-Schmitt-2016 | H1-HindIII-1 | 1 | 0.76 | 0.84 | 0.88 | 0.9 | 0.92 | 0.94 | 0.95 |
| GSE87112-Schmitt-2016 | H1-HindIII-1 | 10 | 0.77 | 0.83 | 0.87 | 0.9 | 0.94 | 0.96 | 0.98 |
| GSE87112-Schmitt-2016 | H1-HindIII-1 | 18 | 0.76 | 0.82 | 0.86 | 0.89 | 0.94 | 0.96 | 0.98 |
| GSE87112-Schmitt-2016 | H1-HindIII-2 | 1 | 0.76 | 0.84 | 0.88 | 0.9 | 0.92 | 0.94 | 0.95 |
| GSE87112-Schmitt-2016 | H1-HindIII-2 | 10 | 0.77 | 0.83 | 0.87 | 0.9 | 0.94 | 0.96 | 0.98 |
| GSE87112-Schmitt-2016 | H1-HindIII-2 | 18 | 0.76 | 0.82 | 0.86 | 0.89 | 0.94 | 0.96 | 0.98 |
| GSE87112-Schmitt-2016 | LV-HindIII-1 | 1 | 0.59 | 0.69 | 0.77 | 0.81 | 0.87 | 0.91 | 0.94 |
| GSE87112-Schmitt-2016 | LV-HindIII-1 | 10 | 0.55 | 0.63 | 0.69 | 0.74 | 0.82 | 0.88 | 0.92 |
| GSE87112-Schmitt-2016 | LV-HindIII-1 | 18 | 0.55 | 0.63 | 0.69 | 0.73 | 0.8 | 0.87 | 0.92 |
| GSE52457-Dixon-2015 | MDC-HindIII-1 | 1 | 0.79 | 0.86 | 0.89 | 0.91 | 0.94 | 0.95 | 0.96 |
| GSE52457-Dixon-2015 | MDC-HindIII-1 | 10 | 0.79 | 0.84 | 0.88 | 0.91 | 0.94 | 0.96 | 0.98 |
| GSE52457-Dixon-2015 | MDC-HindIII-1 | 18 | 0.78 | 0.83 | 0.86 | 0.89 | 0.93 | 0.96 | 0.98 |
| GSE52457-Dixon-2015 | MDC-HindIII-2 | 1 | 0.79 | 0.86 | 0.89 | 0.91 | 0.94 | 0.95 | 0.96 |
| GSE52457-Dixon-2015 | MDC-HindIII-2 | 10 | 0.79 | 0.84 | 0.88 | 0.91 | 0.94 | 0.96 | 0.98 |
| GSE52457-Dixon-2015 | MDC-HindIII-2 | 18 | 0.78 | 0.83 | 0.86 | 0.89 | 0.93 | 0.96 | 0.98 |
| GSE87112-Schmitt-2016 | PO3-HindIII-1 | 1 | 0.51 | 0.64 | 0.73 | 0.79 | 0.86 | 0.91 | 0.94 |
| GSE87112-Schmitt-2016 | PO3-HindIII-1 | 10 | 0.43 | 0.53 | 0.6 | 0.66 | 0.75 | 0.82 | 0.88 |
| GSE87112-Schmitt-2016 | PO3-HindIII-1 | 18 | 0.43 | 0.52 | 0.59 | 0.65 | 0.74 | 0.82 | 0.88 |
| GSE87112-Schmitt-2016 | PA3-HindIII-1 | 1 | 0.53 | 0.65 | 0.73 | 0.79 | 0.86 | 0.9 | 0.93 |
| GSE87112-Schmitt-2016 | PA3-HindIII-1 | 10 | 0.48 | 0.57 | 0.64 | 0.7 | 0.78 | 0.85 | 0.9 |
| GSE87112-Schmitt-2016 | PA3-HindIII-1 | 18 | 0.48 | 0.57 | 0.64 | 0.69 | 0.78 | 0.85 | 0.9 |
